# A ligand/receptor trafficking clock governs self-renewal and abscission dynamics in pluripotent stem cells

**DOI:** 10.64898/2025.12.22.695902

**Authors:** D. Libetti, N. Combémorel, E. Fraimbault, M. Lantelme, X. Gaume, E. Zakiev, Y. Tapponnier, A. Huyghe, G. Furlan, M. Coudurier, L. Claret, C. Rognard, M., S. Akguel, H. Hernandez-Vargas, N. Gadot, F. Forest, M. Koch, I. Aksoy, N. Allegre, C. Chazaud, M. Ramalho-Santos, P. Savatier, E. Bachelard-Cascales, F. Lavial

**Author notes:** Corresponding Author: F. Lavial. These authors contributed equally to this work (D. Libetti, N. Combémorel).

## Abstract

How extracellular cues are temporally integrated to regulate self-renewal and differentiation propensities across the cell cycle remains largely unresolved. We identify a ligand/receptor trafficking clock in rodent and human pluripotent stem cells (PSCs) in which the cyclic turnover of Netrin-1 and its receptors Neo1 and Unc5b (NNU) governs self-renewal capacity and abscission dynamics. In G1, NNU complexes undergo Clathrin-mediated internalization and lysosomal degradation, a process required for timely post-mitotic bridge abscission. At later stages of the cycle, NNU activate Src within early endosomes, inducing a genome-wide redistribution of the transcriptional co-activator Yap1. This reshapes gene regulatory networks by activating stemness- and ectoderm-associated transcriptional programs enriched for Sox2/Nanog binding and by repressing mesodermal- and cell cycle–related targets enriched for Sox2 and Tcf3. Functionally, recombinant Netrin-1 reduces functional heterogeneity and enhances clonogenicity in G1, uncovering a tractable strategy to canalize stem cell behavior. Collectively, our results reveal cell cycle-dependent ligand/receptor trafficking as a temporal clock that directly links membrane dynamics to epigenetic regulation and stem cell fate, opening new avenues for regenerative medicine.

## Introduction

During mammalian development, pluripotent stem cells (PSCs) arise from the inner cell mass of preimplantation blastocysts to form the epiblast, the source of all embryonic lineages (Evans and Kaufman, 1981; Martin, 1981). The fate of PSCs is governed by the integration of multiple signaling pathways—both chemical and mechanical—that collectively maintain pluripotency or initiate differentiation (De Belly et al., 2022). Importantly, PSCs do not simply respond to static levels of individual signals; rather, they integrate dynamic and cumulative signaling inputs over time, as exemplified by fibroblast growth factor/extracellular signal-related kinase (FGF/ERK) and bone morphogenetic protein (BMP) pathways (Hamilton et al., 2019; Teague et al., 2024). This intricate temporal integration enables embryonic stem cells (ESCs) to balance self-renewal with lineage commitment.

*In vitro*, ESCs exhibit heterogeneous responses to differentiation signals, posing challenges for stem cell-based therapies that demand uniform cell populations. Transcriptional heterogeneity contributes to this variability, with mounting evidence implicating the cell cycle machinery as a critical determinant of stem cell responses (Espinosa-Martinez et al., 2024). Notably, the G1 phase represents a restricted window during which PSCs are most responsive to differentiation cues (Pauklin and Vallier, 2013), regulated by factors such as the transcription factor (TF) SMAD2/3 and the chromatin proteins Polycomb and Trithorax (Asenjo et al., 2023; Asenjo et al., 2020; Coronado et al., 2013; Gonzales et al., 2015; Grandy et al., 2016; Pauklin and Vallier, 2013; Sela et al., 2012; Singh et al., 2015). Conversely, S and G2 phases favor maintenance of pluripotency through regulators like p53 and cyclin B1 (Aranda et al., 2023; Gonzales et al., 2015; Neganova et al., 2014). Intriguingly, PSC fate is also linked to the mechanical dynamics of abscission—the final separation of daughter cells after mitosis (Chaigne et al., 2020; Kodba et al., 2025). Together, these findings underscore the importance of coordinated regulation across cell cycle phases and abscission dynamics in controlling PSC identity and behavior. Yet, the molecular mechanisms underpinning this coordination remain largely unknown, with prevailing models overlooking potential roles for signaling pathways. While canonical signaling molecules such as FGF, TGFβ/BMP, and WNT govern ESC biology, their expression level appear constant throughout the cell cycle, suggesting that signal cyclicity does not influence stem cell fate decisions (Dalton, 2015; Singh et al., 2013).

Netrin-1, a laminin-like protein originally characterized as an axon guidance cue, harbor pleiotropic functions that depend on receptor context and include morphogenetic remodeling (Kennedy et al., 1994; Piper et al., 2005). Although Netrin-1 influences embryonic, adult, and cancer stem cells features (Cassier et al., 2023; Furlan et al., 2023; Huyghe et al., 2020; Lengrand et al., 2023; Renders et al., 2021; Sung et al., 2019), the spatiotemporal dynamics of this ligand and its receptors at single-cell resolution, and their mechanisms of signal transduction from membrane to nucleus, remain largely unexplored.

Here, we discovered that the cyclic trafficking of Netrin-1 and its receptors constitutes a previously unappreciated paradigm that dynamically links cell cycle progression, abscission, and Yap1-mediated control of stem cell clonogenicity and differentiation propensities.

## Results

### A Tri-Color Fluorescent Reporter Model Uncovers Single-Cell Heterogeneity of Netrin-1 Ligand and Receptors in Pluripotent Stem Cells

To investigate the single cell-dynamic of the Netrin-1 ligand and its receptors Neo1 and Unc5b (collectively NNU)(Huyghe et al., 2020), we generated a tri-color reporter mouse ESC line (TriC) by inserting fluorescent tags into each gene locus using CRISPR/Cas9 (Figure 1A and Figure S1A). Specifically, the Green fluorescent proteins (GFP), the mTagBFP2 (BFP) and the mCherry (RFP) were inserted into the *Ntn1*, *Neo1* and *Unc5b* loci, respectively, with 2A self-cleaving peptides (Ryan et al., 1991). Following antibiotic selection, we isolated a TriC line with bi-allelic integration of GFP (*Ntn1*) and RFP (*Unc5b*), and mono-allelic integration of BFP (*Neo1*), as measured by genomic PCR and copy number assays (Figure S1B-C). Targeting efficiencies were 40% (*Ntn1*), 83% (*Neo1*) and 77% (*Unc5b*). We next assessed whether the insertions affected gene function or pluripotency. While Neo1 and Unc5b protein levels were unchanged, Ntn1 expression was reduced (Figure S1D). However, this had no strong impact on downstream signaling, as levels of Esrrb and β-catenin remained largely stable (Figure S1D-E)(Huyghe et al., 2020). Furthermore, recombinant Netrin-1 (r-Net) treatment induced Nanog expression in TriC and parental cells to a similar extent, confirming the functionality of the genetically engineered receptors (Figure S1F).

**Figure 1:**
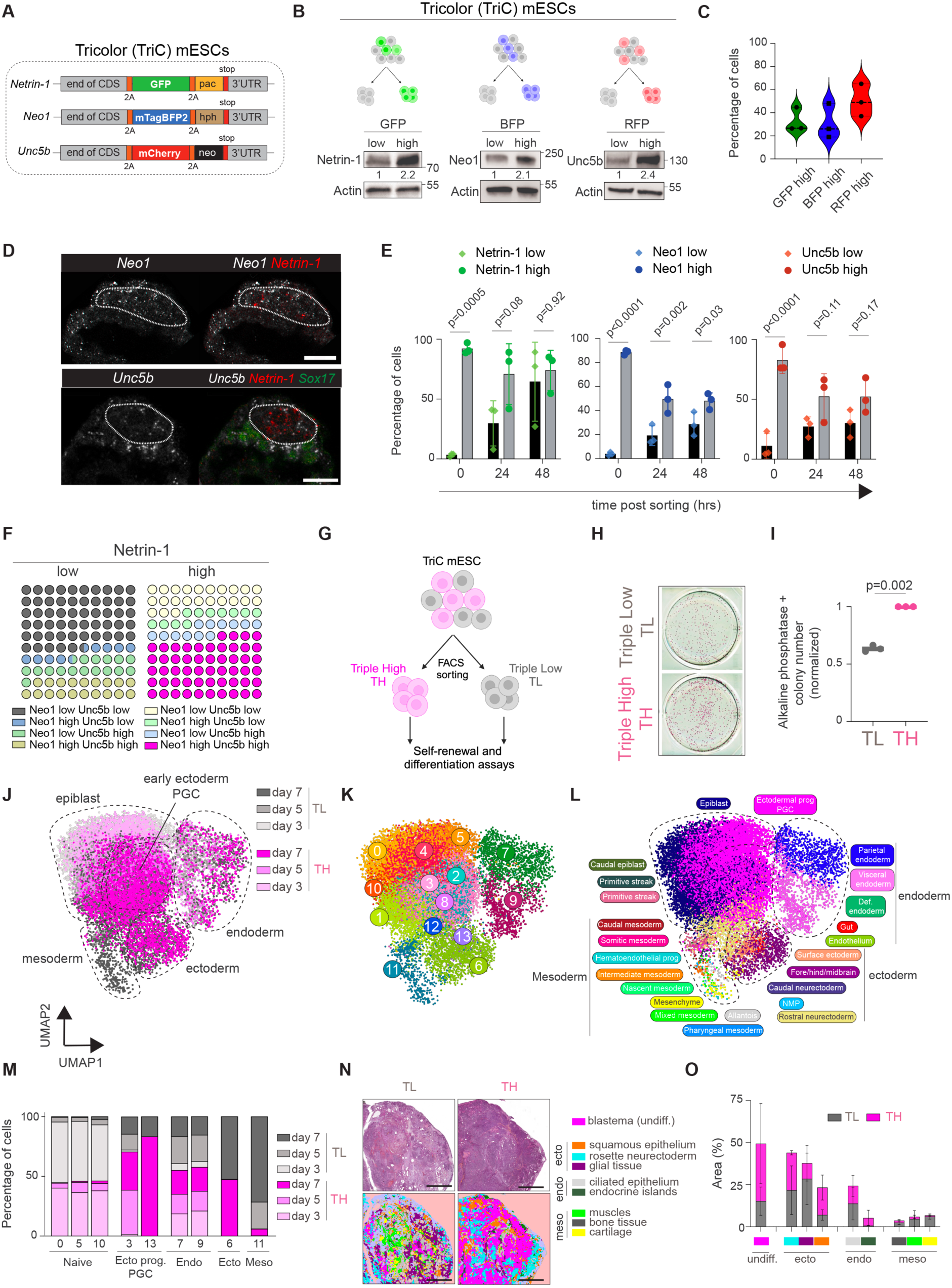
NNU levels discriminates mouse embryonic stem cells (mESCs) with various self-renewal and differentiation abilities. **(A)** Schematic diagram of TriC mESC generation. A triple knock-in fluorescent reporter mESC line was generated by inserting the GFP, mTagBFP2 and mCherry coding sequences after the Ntn1, Neo1 and Unc5b coding sequences. 2A: Self-cleaving peptide; pac: puromycin N-acetyltransferase; hph: hygromycin phosphotransferase; neo: amino 3′-glycosyl phosphotransferase; 3’UTR: 3’ untranslated region. **(B)** Top panel: schematic diagram showing the heterogeneous expression of the fluorescent reporters. Bottom panel: Western blots against Netrin-1, Neo1 and Unc5b proteins in TriC mESC sorted by FACS for the indicated fluorescence. **(C)** Expression of NNU in mouse ESCs. Violin plot representing the percentage of mESCs expressing the indicated factors. ESC: Embryonic stem cells. **(D)** *Netrin-1*, *Neo1* and *Unc5b* localization in pre-implantation embryos. Embryos were fixed around E4.5 and stained for the indicated transcripts. Sox17 is used to mark primitive endoderm cells. The white dotted line highlights the epiblast. **(E)** Bar charts depicting the interconvertibility of mESC subsets expressing different levels of Netrin-1, Neo1 and Unc5b. The percentages of cells were quantified by FACS analysis at the indicated time points. n= 3 independent experiments. Data are the mean +/- sd and two-way ANOVA t-test was used for statistics. **(F)** Relative proportions of mESC subpopulations in culture. The percentage of the indicated subpopulations was quantified by FACS analysis. One experiment representative of three independent experiments is presented. **(G)** Schematic diagram of the procedure used to characterize Triple High (TH) and Triple Low (TL) mESCs. **(H)** Images of colonies stained for alkaline phosphatase (AP). TH and TL cells were FACS sorted and plated in self-renewing conditions for 7 days before staining. **(I)** Corresponding countings of AP+ colonies. n= 3 independent experiments. Data are the mean +/- sd and Student t-test was used for statistics. Value 1 was given to the number of colonies generated by TH cells. **(J-L)** Uniform Manifold Approximation and Projection (UMAP) visualization of scRNA-Seq profiles. TH and TL cells were FACS sorted and subjected to embryoid body formation. The embedding integrates 20,849 pre-processed cells (individual dots), corresponding to three time points of differentiation processed in one sequencing experiment. The cells are coloured by sample (j), by Louvain cluster (k), or by fate class assigned during single-cell reference projection of our data on the atlas of gastrulating embryos (Pijuan-Sala et al., 2019). **(M)** Percentage of cells in each cluster. **(N)** Histological staining of teratomas generated by the injection of FACS-sorted TH or TL cells. Two independent teratomas were analysed per cell line. Scale bars, 2mm. Top panel: brightfield micrographs. Bottom panel: Affiliated structures analysed by Artificial Intelligence. **(O)** Histogram presenting the relative proportion of each differentiated cell type. Data are the mean +/- sd of 3 independent slides.

To assess reporter fidelity, TriC ESCs grown in Serum/LIF were FACS-sorted based on GFP, BFP, or RFP intensity (Figure 1B and S1G). Fluorescence levels correlated with endogenous Netrin-1, Neo1 and Unc5b protein levels, indicating that TriC ESCs constitute a unique opportunity to track NNU dynamics (Figure 1B). Flow cytometry revealed heterogeneous expression of the three fluorescences: ∼32% of TriC ESCs were Netrin-1^high^, 31% Neo1^high^, and 50% Unc5b^high^ under serum/LIF conditions (Figure 1C). Culture in 2i medium significantly increased the proportion of Netrin-1^high^ and Neo1^high^ ESCs (Figure S1H). *Neo1* and *Unc5b* heterogeneity was observed at the transcript level in epiblastic cells of pre-implantation embryos (Figure 1d), in a similar manner as *Ntn1* (Huyghe et al., 2020). Together, these results establish TriC ESCs as a robust model to track NNU signaling at single-cell resolution and reveal inherent heterogeneity in Netrin-1 pathway components both *in vitro* and *in vivo*.

### Netrin-1/Neo1/Unc5b (NNU) levels define mESCs populations with distinct self-renewal and differentiation abilities

We next asked whether mESCs expressing high or low levels of Netrin-1, Neo1, and Unc5b (NNU) differ in self-renewal capacity. Netrin-1^low^ and Netrin-1^high^ mESCs were FACS sorted and replated for colony formation assays. Netrin-1^high^ cells formed significantly more undifferentiated alkaline phosphatase positive colonies than Netrin-1^low^ cells (Figure S2a-b)(Huyghe et al., 2020; Ozmadenci et al., 2015). Similar results were observed comparing Neo1^high^ vs Neo1^low^ and Unc5b^high^ vs Unc5b^low^ subpopulations (Figure S2A-B). When replated and analyzed over 48 hours, all six subpopulations dynamically interconverted, returning to an equilibrium distribution, indicating these states are reversible (Figure 1E).

Detailed FACS profiling revealed eight subpopulations based on NNU expression, with the dominant state being Triple High (TH; Netrin-1^high^/Neo1^high^/Unc5b^high^; 54% of Netrin-1^high^ cells) and Triple Low (TL; Netrin-1^low^/Neo1^low^/Unc5b^low^; 56% of Netrin-1^low^ cells) (Figure 1F). These findings prompted us to conduct a multi-scale characterization of TH and TL mESCs (Figure 1G). Transcription factors previously linked to ESC heterogeneity (Nanog, Esrrb, Hes1, Rex1)(Chambers et al., 2007; Kobayashi et al., 2009; Kumar et al., 2014; Torres-Padilla and Chambers, 2014) were similarly expressed in TH and TL cells, suggesting these subpopulations are distinct from known subsets (Figure S2C). Functionally, TL cells exhibited reduced colony-forming efficiency compared to TH cells despite slightly higher Oct4 levels (Figure 1H-I and S2D).

To assess differentiation bias, embryoid bodies (EBs) derived from TL and TH cells were analyzed over 3, 5, and 7 days. RT-qPCR showed that TL-derived EBs expressed higher mesodermal markers (*Brachyury*, *Mixl1*), whereas TH-derived EBs were enriched for early ectodermal and primordial germ cell markers (*Rhox9*, *Dppa3*) at day 7, with no significant differences in endodermal markers (Figure S2E). Single-cell RNA sequencing was next conducted in similar settings. After preprocessing 20,859 cells, k-nearest neighbors (k-NN) clustering on the dimensionality reduction embedding of principal components analysis (PCA), further resolved 14 clusters corresponding to germ layer derivatives (Figure 1J-K and S2F-G). Reference mapping based on an atlas of gastrulating embryos (Pijuan-Sala et al., 2019) further refined the affiliation of the clusters to embryonic germ layers (Figure 1L and Figure S2H). Consistent with bulk data, TL-derived cells predominated in mesodermal clusters (e.g., cluster 11: *Brachyury*, *Cdx2*), while TH-derived cells were enriched in ectodermal and PGC clusters (clusters 3, 13: *Rhox6*/*Rhox9*/*Dppa3*) (Figure 1M and S2G-H). TH and TL cells equally contributed to the epiblastic (0, 5 and 10) and to the endodermal clusters (7 and 9: *Gata4, Gata6* and *Sox17)* (Figure 1M and S2G-H). *In vivo*, teratomas formed from sorted TL and TH cells maintained at least partially these lineage biases (Figure 1N-O). The HALO Artificial intelligence tool was trained by pathologists to specifically affiliate and quantify histological structures. Interestingly, TL-derived teratoma contained more mesodermal derivatives while TH-derived tissues were enriched in structures maintaining a globally undifferentiated or early ectodermal morphology (Figure 1N-O). Together, these findings reveal that mESCs with high or low NNU expression define functionally distinct, interconvertible subpopulations with divergent self-renewal and lineage differentiation potentials.

### NNU Expression Correlates with Chromatin Accessibility and Cell Cycle–Associated Transcriptional Programs

To explore the molecular basis underlying the functional differences between TL and TH mESCs, we performed integrated RNA-seq and ATAC-seq on FACS-sorted populations. TH cells displayed a greater number of ATAC-seq peaks, indicating a globally more accessible chromatin landscape (Figure S3A). While TL cells had more promoter-associated peaks, TH cells showed increased accessibility in intronic regions (Figure S3B). Consistent with their lineage biases, chromatin accessibility was elevated at ectodermal loci (e.g., Sox1) in TH cells and at mesodermal loci (e.g., Tbx15) in TL cells (Figure 2A). Notably, these changes occurred without corresponding differences in transcript levels (Figure 2B), indicating that NNU dynamic is associated with chromatin changes that may precede transcriptional regulation. RNA-seq identified 477 differentially expressed genes (DEGs; adj. p < 0.05, |log₂FC| > 1), with 367 upregulated in TH cells (Figure 2C). Gene Set Variation Analysis (GSVA) using these 367 genes as a “TH score” confirmed a strong dependency on NNU: the score was reduced in Ntn1, Neo1, and Unc5b CRISPR/Cas9 knockout and elevated in Netrin-1 Dox-inducible mESCs (Figure 2D). Gene Ontology (GO) analysis unexpectedly linked these genes to “mitotic nuclear division” and “endocytosis,” suggesting a connection to cell cycle regulation (Figure 2E).

**Figure 2:**
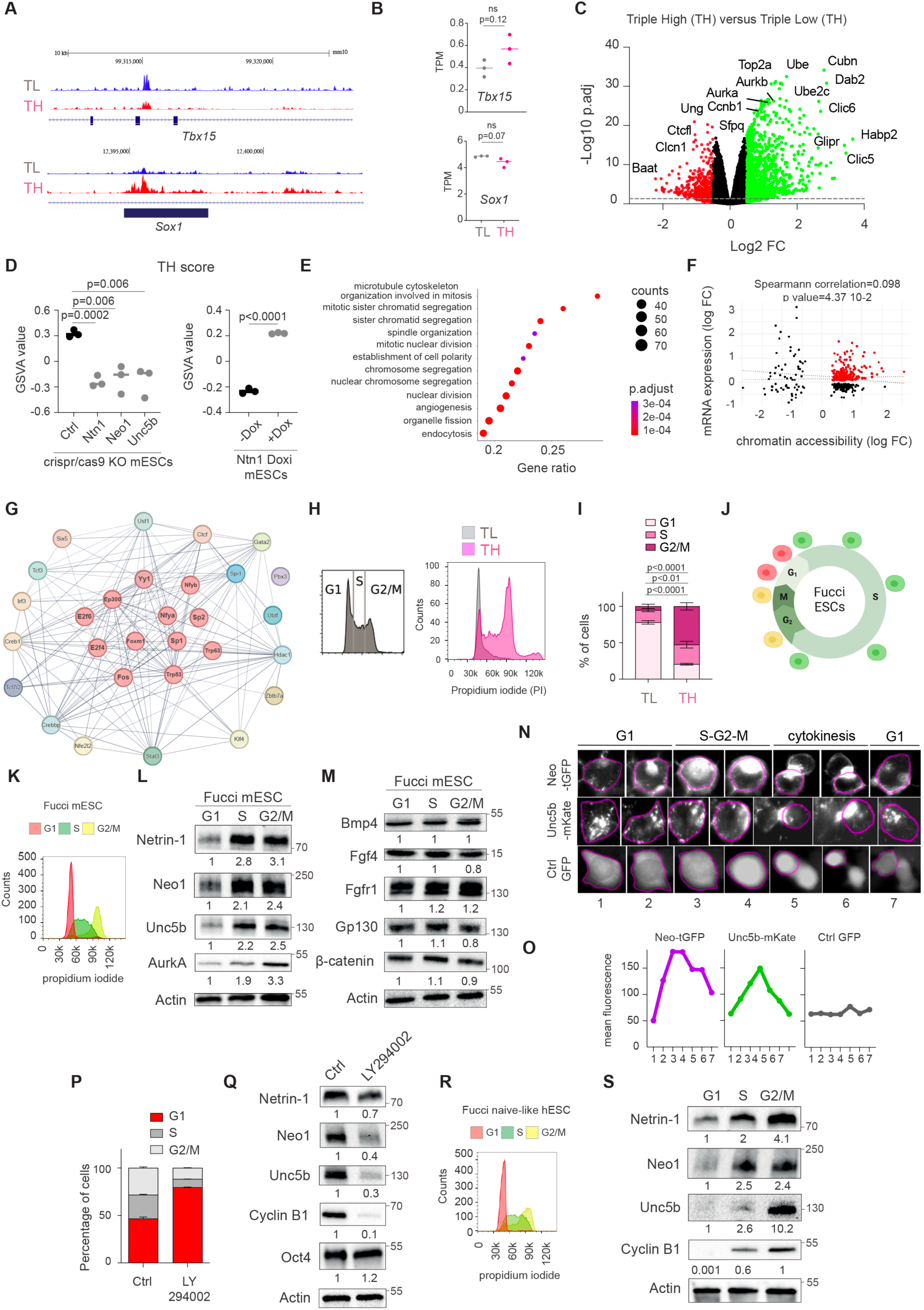
NNU level discriminate mESCs with different epitranscriptomic features connected to cell cycle regulators. (**A**) Example of ATAC chromatin sites at the Tbx15 and Sox1 loci in TH and TL cells. (**B**) Graphs depicting *Tbx15* and *Sox1* transcript level in TH and TL cells. TPM: transcripts per kilobase million. n= 3 independent experiments. Data are the mean +/- sd and Student t-test was used for statistics. (**C**) Volcano plot showing transcripts in Triple High (TH) versus Triple Low (TL) cells. Each dot corresponds to a transcript. The single horizontal dotted line corresponds to a threshold of adjusted p-value of 0.05. Green dots are genes up-regulated with a log2[FC]>0.5 and red dots correspond to down-regulated genes with log2[FC]<-0.5. (**D**) Patterns of TH score. A TH score was defined (see main text for detailed description). Gene Set Variation Analysis (GSVA) was conducted on the transcriptomes of the indicated cells, and statistical significance was evaluated with the Student’s t-test. (**E**) Gene Set Enrichment Analysis (GSEA) on differentially expressed genes between TH and TL cells against Gene Ontology (Biological Process) pathways. P-values were adjusted using Benjamini–Hochberg correction. (**F**) Starburst plot showing the correlation between changes in gene expression and chromatin accessibility in TH and TL cells. See main text for details. (**G**) Prediction of master regulators of TH and TL cells. A list of 217 genes, upregulated and more accessible in TH cells, was extracted from and tested for ORA of ENCODE/ChEA consensus transcription factor (TF) terms. The top 25 factors are presented using String(Szklarczyk et al., 2023). (**H**) Representative cell cycle profile of TriC mESCs. Left panel: global TriC mESC population; right panel: FACS sorted TH and TL cells. Cell cycle analysis was conducted on TriC ESCs by FACS using propidium iodide. (**I**) Quantification of cells in (h). n= 3 independent experiments. Data are the mean +/- sd and two-way ANOVA-test was used for statistics. (**J**) Diagram depicting the fluorescence of Fucci mESCs during the cell cycle. (**K**) Representative cell cycle profile of Fucci mESCs. Cells were FACS sorted in G1, S and G2/M and analyzed for cell cycle using propidium iodide to demonstrate the efficacy of the Fucci system. (**L-M**) Western blot for the indicated proteins conducted on FACS sorted Fucci mESCs. (**N**) Representative images of cell cycle phases in Neo-TurboGFP, Unc5b-mKate vs Ctrl TurboGFP cell lines (numbered 1 to 8). The magenta circle marks the membrane of a single cell on time-lapse imaging. (**O**) Quantification of mean membrane fluorescence intensity for each picture. (**P**) Percentage of cells in each phase of the cell cycle in mESCs after treatment with LY294002 for 24h. Data correspond to one experiment representative of 2 independent experiments. (**Q**) Western blot for samples corresponding to (P). (**R**) Representative cell cycle profile of Fucci naïve-like hESC. Cells were FACS sorted in G1, S and G2/M and analysed for cell cycle using propidium iodide to demonstrate the efficacy of the Fucci system. (**S**) Western blot for the indicated proteins.

To integrate transcriptional and epigenetic data, we generated a starburst plot highlighting 217 genes with both increased expression and chromatin accessibility in TH cells (FDR < 0.05; Figure 2F). Over-representation analysis (ORA) of ENCODE/ChEA TF binding data revealed significant enrichment for cell cycle–associated transcription factors, including p53, E2F4, E2F6, Sp1, Sp2, NFYA/B, YY1, p300, and FoxM1 (Figure 2G and S3C)(Fischer, 2017). Together, these findings reveal that NNU expression dynamics are closely coupled to widespread transcriptional and epigenetic changes and uncover a previously unrecognized link between Netrin-1 signaling and cell cycle in pluripotent stem cells.

### NNU expression is cyclic in mouse and human ESCs

To further explore the relationship between NNU signaling and the cell cycle, we performed Gene Set Variation Analysis (GSVA) on TL and TH cells using established G1/S and G2/M gene signatures. TL cells were enriched for G1/S-phase genes, whereas TH cells displayed a strong G2/M signature (Figure S3D). Cell cycle profiling with propidium iodide confirmed this distribution: 78% of TL cells were in G1, while 80% of TH cells were in G2/M (Figure 2H-I), a pattern also observed when sorting cells based on individual GFP, BFP, or RFP levels (Figure S3E). Unlike canonical signaling molecules such as FGF, TGFβ/BMP, and WNT—whose activity remains constant throughout the cell cycle (Dalton, 2015) - we hypothesized that NNU expression may be cyclic. To test this, we used mESCs expressing the FUCCI reporter, which labels G1 (RFP⁺), S (GFP⁺), and G2/M (RFP⁺/GFP⁺) cells via the phase-specific degradation of Cdt1 and Geminin (Sakaue-Sawano et al., 2017) (Figure 2J-K). FACS analysis yielded 18.1% G1, 58.7% S, and 23.2% G2/M cells, confirmed by propidium iodide staining (Figure S3F). RNA-seq and GSVA validated the accuracy of cell cycle assignment (Figure S3G). Western blotting revealed that Netrin-1, Neo1, and Unc5b were strongly downregulated in G1, alongside low Aurora kinase A levels, consistent with prior reports (Figure 2L)(Naso et al., 2021). This cyclicity appeared mainly specific to NNU, as core components of other signaling pathways (e.g., Gp130, Bmp4, β-catenin, Fgf4, Fgfr1) remained constant across cell cycle phases (Figure 2M). Importantly, transcript levels of *Ntn1*, *Neo1*, and *Unc5b* did not vary significantly, pointing to the prevalence of post-transcriptional regulations (Figure S3H).

To directly monitor receptor membrane dynamics, we tracked Neo1-TurboGFP and Unc5b-mKate fusions expressed under a constitutive promoter in polyclonal mESC lines. Unlike unfused TurboGFP control, both fusion proteins exhibited cyclic membrane accumulation, peaking during S/G2/M and diminishing in G1 (Figure 2N-O). Pharmacological enrichment of G1-phase cells using the PI3K inhibitor LY294002 (Jirmanova et al., 2002; Wang et al., 2017) led to a marked decrease in NNU protein levels, mirroring the reduction in Cyclin B1, a known S–G2/M marker (Figure 2P-Q)(Hayward et al., 2019). Conversely, synchronizing cells in mitosis using demecolcine (Jirmanova et al., 2002), that blocks mitotic spindle formation, increased NNU expression, which declined progressively upon release as cells transitioned into G1 (Figure S3I-K).

We next asked whether this cyclic expression is conserved in human ESCs. FUCCI primed hESCs were converted to a naïve-like state using PD0325901, XAV939, Gö6983 and LIF (Bredenkamp et al., 2019). Flow cytometry and propidium iodide analysis confirmed efficient cell cycle staging (Figure 2R). Western blotting revealed strong downregulation of Netrin-1, Neo1, and Unc5b in G1, mirroring the pattern in mESCs (Figure 2S). While some degree of cyclicity was retained in primed hESCs, especially for Unc5b and Netrin-1, the amplitude was reduced (Figure S3L-M). In both naïve and primed hESCs, NNU transcript levels remained stable across cell cycle phases (Figure S3N-O), again indicating the existence of post-transcriptional controls. Together, these results establish Netrin-1 and its receptors cyclicity as a hallmark of mouse and human naïve pluripotency.

### Clathrin-mediated endocytosis and lysosomal degradation contribute to NNU protein cyclicity in mouse and human ESCs

Given that *Ntn1*, *Neo1*, and *Unc5b* transcript levels remain stable across the cell cycle, we investigated post-transcriptional mechanisms that might regulate their protein-level cyclicity. To determine whether translational regulation contributes to NNU dynamics, we performed polysome profiling on FACS-sorted Fucci mESCs. While *AurkB* mRNA was depleted from heavy polysomes in G1—consistent with low protein expression—*Ntn1*, *Neo1*, and *Unc5b* transcripts showed no significant differences in polysome loading across G1, S, or G2/M phases (Figure S4A-B). This data rules out translation efficiency as a primary driver of NNU periodicity.

As several receptors from the nervous system are known to be internalized and degraded in endolysosomes (Harrington et al., 2011), we first asked whether fluctuations in global endocytosis levels during the cell cycle could contribute to NNU cyclicity. Even if endocytosis globally increased in differentiated mESCs (De Belly et al., 2021), whether it fluctuates during the cell cycle in naïve stem cells remains unknown. Using dextran uptake assays on Fucci-sorted mESCs, we found significantly more mESCs highly incorporating Dextran (Dextran^High^) in G1 compared to S/G2/M, as quantified by FACS and single-cell imaging (Figure 3A-C). This cell cycle–dependent fluctuation was also observed in naïve-like hESCs (Figure 3D-E). Notably, this high endocytic activity in G1 cells occurred without corresponding changes in FGF activity, as Fgf4 and Fgfr1 levels remained constant during the cell cycle (Figure 2M), indicating that it is mainly uncoupled from the events occurring in differentiated cells (De Belly et al., 2021).

**Figure 3:**
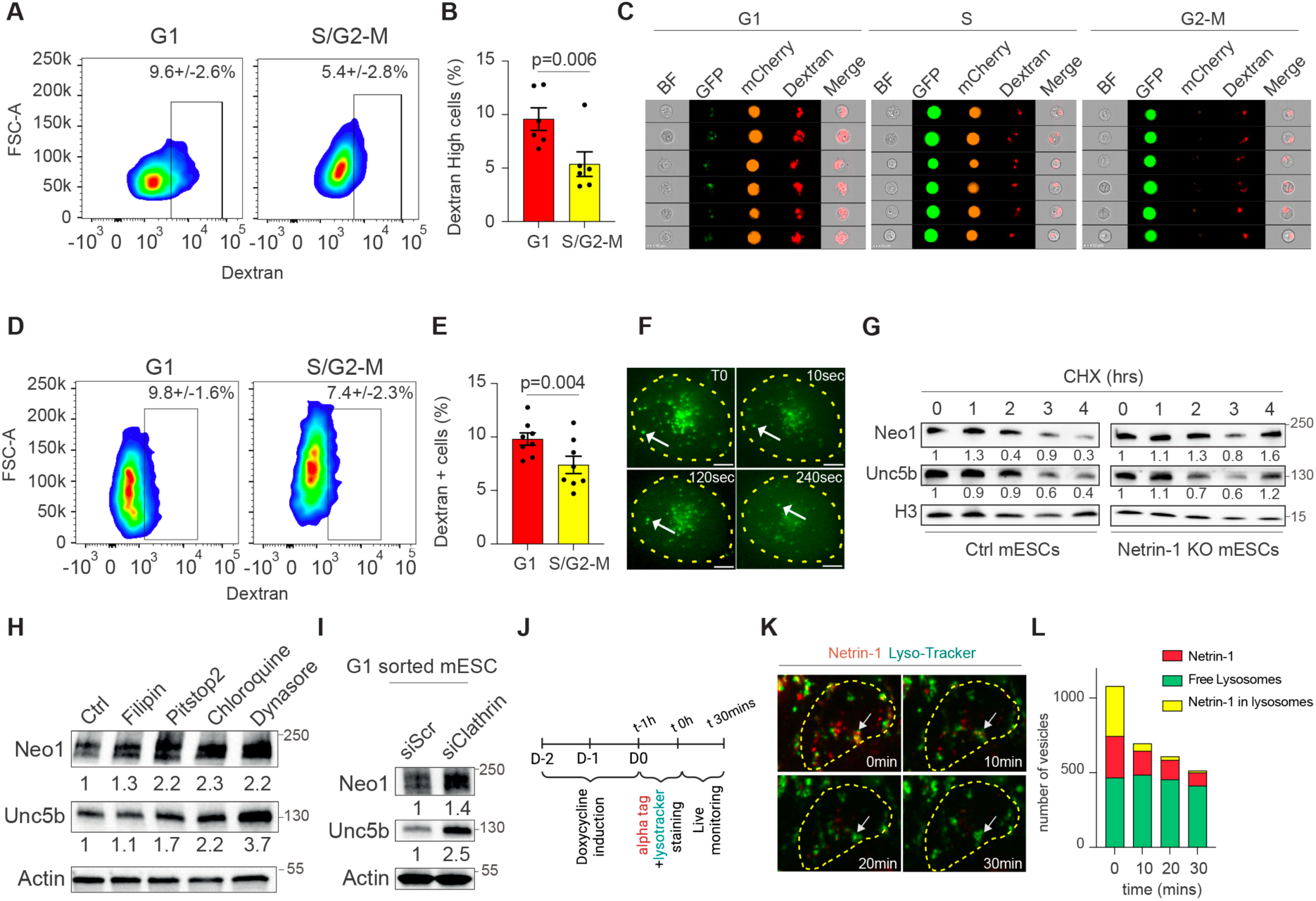
Clathrin-mediated endocytosis and lysosomal degradation regulate NNU levels during the cell cycle. (**A**) Representative FACS profile of Fucci mESCs incorporating Dextran. Fucci mESCs were incubated with Dextran for 30 mins (see Methods for details) and analyzed by FACS. (**B**) Bar plot quantifying Dextran^high^ cells in G1 and S/G2-M Fucci mESCs. n= 6 independent experiments. Data are the mean +/- sd and Student t-test was used for statistics. (**C**) Representative pictures of Fucci mESCs incorporating Dextran. Brightfield, GFP, mCherry and Dextran (coupled to the AF647) imaging of Fucci cells in G1 (left panel), S (middle panel) and G2-M phases (right panel). Magnification x40. (**D**) Representative FACS profile of Fucci naïve-like hESCs incorporating Dextran. (**E**) Bar plot quantifying Dextran^high^ cells in G1 and S/G2-M Fucci naïve-like hESCs. n= 8 independent experiments. Data are the mean +/- sd and Student t-test was used for statistics. (**F**) Time-lapse imaging of Neo1–TurboGFP, acquired at one frame per second, showing intracellular trafficking. The white arrow traces the trajectory of a single internalized protein, initially indicated by the white arrow. (**G**) Quantification of Netrin-1, Neo1 and Unc5b protein stability during cycloheximide treatment for the indicated times. n= 3 independent experiments. Data are the mean +/- sd. (**H**) Western blot against Neo1 and Unc5b in mESCs treated with the indicated molecules for 24h. (**I**) Western blot against Neo1 and Unc5b in Fucci mESCs FACS sorted in G1. Cells were transfected with siRNAs 48h prior to FACS sorting. (**J**) Schematic representation of experimental design to track Netrin α-tagged internalization and fusion with lysosome on alive cells over time. (**K**) Microscopy image showing vesicles containing α-tagged Netrin-1(red fluorescence) and lysosomes (green fluorescence). The white arrow indicates a co-staining site showing the degradation of α-Tag Netrin-1, and the yellow dashed line delineates the contour of a single cell. (**L**) Bar plot showing the quantification of green (lysotracker), red (Netrin-alpha-tag) or co-labelling fluorescent vesicles as a function of time in minutes. Quantification was performed on two independent recordings of a monolayer culture (total co-stained vesicles counted at t0: 524).

We next investigated more specifically the membrane dynamics of Neo1 and Unc5b in mESCs. We first found that Neo1–TurboGFP exhibited dynamic intracellular trafficking in live mESCs (Figure 3F and Supplementary Movie 1). To characterize Neo1/Unc5b protein turnover, we first blocked protein synthesis using cycloheximide. The two receptors had short half-lives (∼2 hours), which were extended in Ntn1 knockout cells—revealing a ligand-dependent effect on receptor stability (Figure 3G and Figure S4C-D). We next dissected pharmacologically the endocytic pathways potentially involved. Inhibition of caveolae/raft-mediated endocytosis (Filipin III) had no significant effect, while Clathrin-mediated endocytosis blockade (Pitstop2, Dynasore), together with inhibition of lysosomal degradation (Chloroquine), led to significant accumulation of Neo1/Unc5b (Figure 3H). These findings indicate that Neo1 and Unc5b undergo Clathrin-dependent internalization and lysosomal degradation in mESCs. Importantly, RNAi-mediated knockdown of Clathrin led to a significant accumulation of Neo1 and Unc5b in FACS-sorted G1 mESCs, indicating that Clathrin-dependent endocytosis of Neo1 and Unc5b in G1 contributes to their degradation and cyclicity (Figure 3I).

We next assessed more precisely whether the ligand Netrin-1 is internalized in endosomes but also degraded in lysosomes. To do so, we generated a doxycycline-inducible mESC line that allows to monitor the intracellular route of newly synthesized α-tagged Netrin-1 (Figure 3J). After 48 hours of doxycycline induction, we combinatorially labelled α-tagged Netrin-1 and lysosomes with Lysotracker. Extracellular Netrin-1 was rapidly incorporated into lysosomes and subsequently degraded (Figure 3K-L, Supplementary Movie 2). Together, these results show that cyclic NNU downregulation in G1 is driven by elevated Clathrin-mediated endocytosis and subsequent lysosomal degradation.

### Endocytosis of Neo1 and Unc5b Is Required for coordinated Abscission in mESCs

While tracking Neo1–TurboGFP and Unc5b–mKate dynamics, we observed their accumulation at the intercellular bridge between daughter cells (Supplementary Movies 3–4). Additionally, the cell cycle distribution of Neo1-TurboGFP and Unc5b-mKate mESCs revealed an accumulation of cells in G2/M when compared with control (Figure 4B-C and Figure S4E-F). Previous studies have shown that abscission is tightly regulated in naïve mESCs, which maintain stable intercellular bridges in G1 (Chaigne et al., 2020). Given the endocytic behavior of Neo1 and Unc5b, we hypothesized that their clearance may be required for proper bridge resolution. To test this, we generated doxycycline-inducible mESC lines for tagged versions of Unc5b or Neo1 to enable exogenous expression in G1 (Figure 4d). Functional validation of inducible Unc5b confirmed ligand responsiveness and effective immunoprecipitation *via* Strep-tagIII (Figure 4E and S4G-H). We found that Neo1 and Netrin1 were co-immunoprecipitated along Unc5b, indicating our capability to analyse the interactome of the NNU complex. To probe the composition of this complex, we performed affinity purification followed by mass spectrometry (AP-MS). Among 115 interactors, we identified proteins involved in receptor synthesis, trafficking, and—critically—regulators of cytokinesis and abscission (Figure 4F-H, Supplementary Table 1). These included Anillin (contractile ring stabilization), Septin2 (microtubule scaffolding), and TSG101 (ESCRT recruitment at the midbody), suggesting that the NNU complex might be functionally linked to the abscission machinery. In line with this, exogenous expression of Neo1 or Unc5b led to reduced proliferation (Figure S4I-J), prompting us to directly assess abscission dynamics. α-Tubulin immunostaining revealed a significantly higher number of intercellular bridges in Neo1- and Unc5b-overexpressing cells, indicative of delayed abscission (Figure 4I-J and S4K-L). Notably, chemical inhibition of Clathrin-mediated endocytosis (Pitstop2) phenocopied this effect, further supporting a model in which receptor internalization is required for bridge resolution (Figure 4K). Together, these findings reveal an unexpected role for Neo1 and Unc5b in coordinating abscission. Their endocytosis during G1 ensures timely clearance, thereby enabling proper daughter cell separation in mESCs.

**Figure 4:**
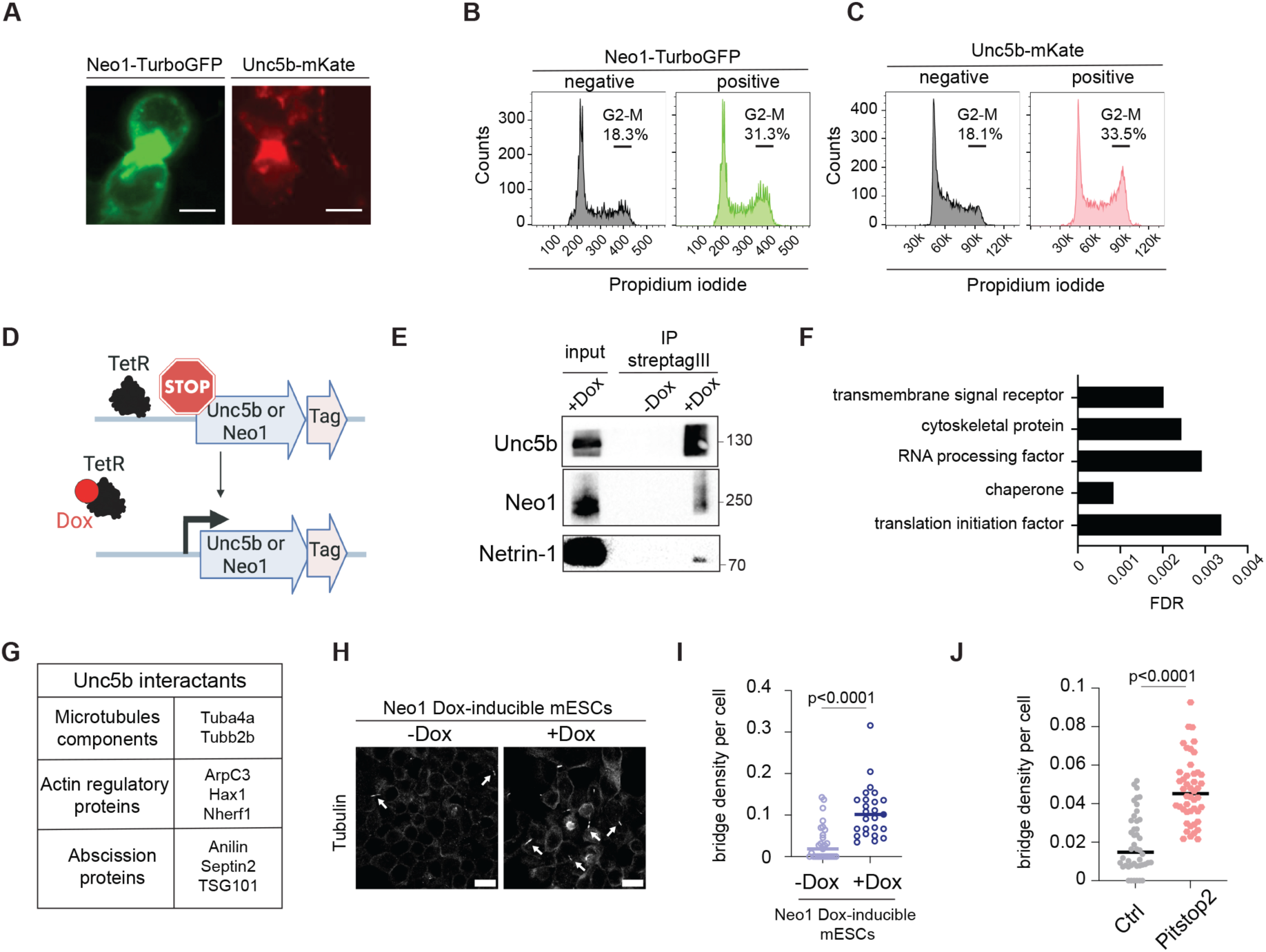
Neo1/Unc5b endocytosis is required for proper bridge abscission. (**A**) Representative example of dividing mESCs expressing fluorescent reporters for Neo1 or Unc5b with white arrowheads showing the fluorescence accumulation between the cells. Left panel: mESC expressing Neo1-TurboGFP Right panel: mESC expressing Unc5b-mKate ESC: Embryonic stem cell. (**B-C**) Representative cell cycle profile of mESCs expressing low and high levels of fluorescent reporters for Neo1 or Unc5b. Left panel: mESC expressing Neo1-TurboGFP Right panel: mESC expressing Unc5b-mKate ESC: Embryonic stem cell. Cell cycle analysis was conducted on mESCs by FACS using propidium iodide. (**D**) Schematic diagram of the experimental design. (**E**) Western blot for Unc5b, Neo1 and Netrin1 after immunoprecipitation of StreptagIII. (**F**) Panther protein class of interactants of Unc5b-StreptagIII. FDR: False Discovery Rate. (**G**) Table of interactants of Unc5b-StreptagIII related to cytoskeleton. (**H**) Representative confocal image of mESCs expressing exogenously Neo1 stained for α-tubulin (white). Left panel: non treated Right panel: doxycycline treatment for 14 days. A maximum Z projection is shown. Scale bar: 20 µm. (**I**) Dot plot showing the fraction of cells with tubulin bridges (number of bridges divided by number of cells in a given analysis frame) in Neo1 Dox-inducible mESCs non treated or treated with doxycycline for 14 days. Mean is shown. n = 3 independent experiments. Student T test was used. **(J)** Dot plot showing the fraction of cells with tubulin bridges (number of bridges divided by number of cells in a given analysis frame) in WT mESCs non treated or treated with Pitstop2 for 48h. Mean is shown. n = 3 independent experiments. Student T test was used.

### NNU Activate a Src–Yap1 Signaling Axis in mESCs via Endocytosis

To elucidate how Netrin-1 and its receptors (NNU) transduce signals during the cell cycle, we investigated pathways beyond WNT and MAPK, whose activities remain constant across cell cycle phases in mESCs (Huyghe et al., 2020). Given the central role of kinases in signal transduction, we performed kinome profiling using peptide arrays (196 phosphotyrosine and 144 phosphoserine/threonine substrates) on lysates from mESCs treated for 24 hours with vehicle or recombinant Netrin-1 (r-Net) (Figure 5A, Supplementary Table 1). Analysis of phosphorylation patterns identified several candidate kinases modulated by r-Net, notably 11 members of the Src family, including Src itself. Immunoblotting confirmed Src activation upon r-Net treatment, as indicated by increased phosphorylation at Tyr416 (Y416) (Figure S5A). Src has been previously shown to activate the Hippo pathway effector Yap1 (Benham-Pyle et al., 2015; Guillermin et al., 2021; Taniguchi et al., 2015). We therefore hypothesized that NNU-mediated Src activation might regulate Yap1. Using gain- and loss-of-function approaches—including r-Net stimulation and NNU knockout mESCs—we found that Netrin-1 induces phosphorylation of Yap1 at Tyr357 (Y357), a modification associated with transcriptional activation, and promotes dephosphorylation at Ser127 (S127), which relieves cytoplasmic sequestration (Figure 5B-C and S5B).

**Figure 5:**
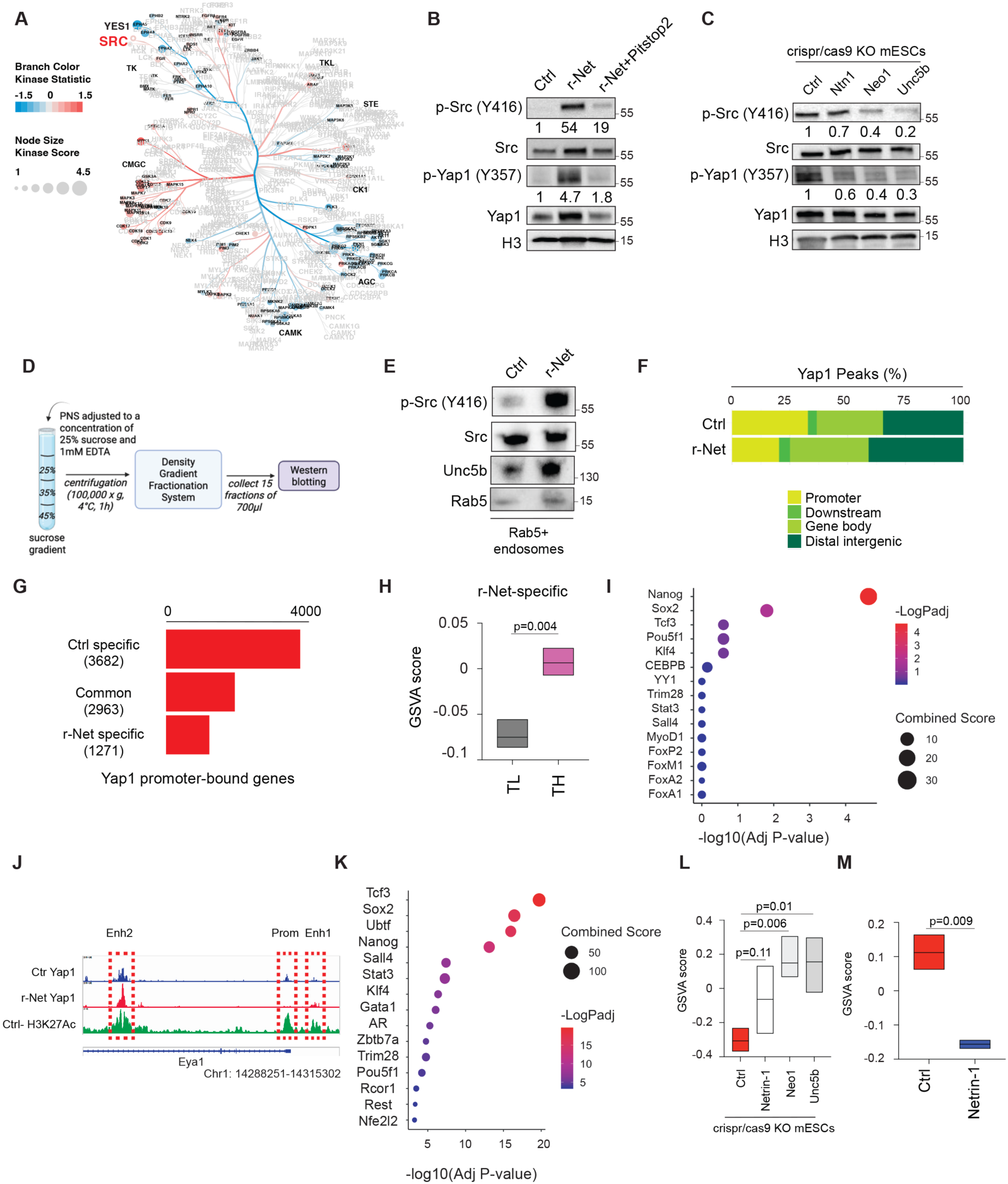
Clathrin-mediated endocytosis of NNU activate a Src/Yap1 signalling axis in ESCs. (**A**) Kinome activity profiling of protein lysates from mESCs treated for 24 hours with vehicle or recombinant Netrin-1. (**B**) Western Blot for phosphorylated forms of Src on Tyr416 and Yap1 on Tyr357 in WT mESCs in non-treated and following recombinant Netrin-1 treatment, with or without the Clathrin inhibitor Pitstop2. (**C**) Western blot for the indicated proteins conducted on Crispr/cas9 KO mESCs. (**D**) Scheme showing the isolation and purification of endosomal fractions on a sucrose gradient. (**E**) Western Blot for phosphorylated Src on Tyr416 on early endosomes fractions of mESC treated or not with rNet for 30 minutes. (**F**) Global distribution of Yap1 peaks in mESCs, Ctrl or treated with recombinant Netrin-1 for 24 hours. (**G**) Number of genes presenting Yap1 binding on their promoter region. (**H**) GSVA score for genes newly bound by Yap1 following r-Net treatment in TH and TL cells expressing high or low endogenous levels of NNU, respectively. (**I**) EnrichR Score for the genes from (H). (**J**) IGV visualization of the Eya1 locus for Yap1 binding and H3K27Ac signal. (**K**) EnrichR Score for the genes showing reallocation of Yap1 from promoter to enhancer. (**L-M**) GSVA score for the genes from (K) genes in the indicated cell lines. On the right panel, Netrin-1 refers to Netrin-1 Dox-inducible mESCs treated with Dox for 24 hours. Student T-test was used.

We showed next that the activation of this NNU/Src/Yap1 axis mainly occurs *via* endocytic trafficking, as its chemical inhibition (Pitstop2) decreases Src/Yap1 activation (Figure 5B)(Huyghe et al., 2020). In line with this, we purified Rab5⁺ early endosomes using a sucrose density gradient (25–45%)(Walker et al., 2016) and found that both Unc5b and phosphorylated Src co-localize in these compartments (Figure 5D-E and S5C), reinforcing the notion that endosomes serve as signaling hubs for NNU. Collectively, these findings demonstrate that the internalization of Netrin-1 and its receptors into Rab5⁺ endosomes facilitates Src activation and downstream Yap1 signaling, thereby linking endocytosis to cell cycle–dependent signal transduction.

### NNU reconfigure Yap1 binding on the chromatin

Yap1 functions as a transcriptional co-regulator, exerting its activity through DNA-binding partners such as Tead1–4 and AP-1 (Fos–Jun dimers)(Zanconato et al., 2015). To assess how NNU influence Yap1 genomic occupancy, we performed chromatin immunoprecipitation followed by sequencing (ChIP-seq) in mESCs treated with recombinant Netrin-1 (r-Net) or vehicle control. Overall, r-Net treatment did not significantly alter the number of Yap1 peaks or associated genes (20,942 peaks/13,175 genes in control vs. 21,211 peaks/12,608 genes in r-Net). As expected, MEME-ChIP motif analysis identified Tead and Yy1 as primary binding motifs (Figure S5D)(Sun et al., 2020). However, a notable redistribution of Yap1 binding sites across genomic features was observed following r-Net treatment. Specifically, the number of promoter-bound peaks decreased by 37% (from 6,871 to 4,318), while binding in downstream, intragenic, and distal intergenic regions increased by 20%, 19%, and 18%, respectively (Figure 5F).

We next conducted a qualitative analysis of Yap1 binding on promoters. Netrin-1 induced both loss (3,682 genes) and gain (1,271 genes) of Yap1 occupancy, while binding on 2,963 genes remained unchanged (Figure 5G). Gene Set Variation Analysis (GSVA) revealed that Yap1-bound genes in response to r-Net were significantly more expressed in TH cells (enriched in S/G2-M) when compared to TL cells (enriched in G1), consistent with the higher NNU activity during S/G2/M (Figure 5H). EnrichR analysis further demonstrated that these loci are enriched for Nanog and Sox2 binding in mESCs (Figure 5I), suggesting that Netrin-1 facilitates Yap1 recruitment to Nanog/Sox2-target genes that are highly expressed in S/G2/M.

While Yap1 is traditionally associated with promoter-driven gene activation, recent studies indicate that it also modulates transcription via enhancer regions, often through long-range chromatin interactions (Cebola et al., 2015; Zhu et al., 2019). Cross-referencing our ChIP-seq data with published H3K27ac profiles (Chronis et al., 2017) revealed that 67% (10,918/16,541) of Yap1 non-promoter peaks in mESCs correspond to active enhancers. Notably, in 1,052 genes, we noticed that r-Net treatment induces a switch in Yap1 occupancy from the promoter to enhancer region(s) of the same gene (Figure 5J). These genes were significantly associated with Sox2 and Tcf3 binding (Figure 5K). Exploration of NNU gain and loss-of-function RNA-seq datasets demonstrated that they globally repress the expression of these genes (Figure 5L-M). Gene set enrichment analysis using Pantherdb highlighted genes related to differentiation in embryonic- or extra-embryonic layers (*Gata6*, *Sox4*, *Tfap2c*) together with gastrulation/primed pluripotency (*Nf2*, *Apln*, *Nodal*, *Tcf15*)(Mi et al., 2019; Thomas et al., 2022). Genes related to the cell cycle transition (*Lmnb1*, *Ccnd1*, *Cdk1*, *Rad21*, *Ccne1*, *Cks2*, *Cep120*) but also Actin cytoskeleton (*Chmp4c*, *Capza2*, *Cdc42ep1*, *Pkd1*, *Cdh5*, *Tpm1*) were also highlighted. Collectively, this data revealed that Netrin-1 triggers a significant redistribution of Yap1 on the genome, including (1) Yap1 recruitment to the promoters of Sox2/Nanog-bound genes that are upregulated in S/G2/M and (2) Yap1 reallocation from promoters to enhancers of Sox2/Tcf3-target genes that are transcriptionally downregulated by NNU.

### Modulation of NNU Activity Alters the Transcriptomes of Mouse and Human Pluripotent ESCs

To investigate how NNU periodicity influences gene expression programs in ESCs, we first used a complementation strategy involving recombinant Netrin-1 (r-Net) treatment of G1 cells. Importantly, we showed first that r-Net treatment was sufficient to induce the expression of its cognate receptors Neo1 and Unc5b in G1 FACS sorted cells (Figure S5E), enabling us to assess downstream transcriptional responses. We performed RNA-seq on G1 FACS-sorted mESCs treated or not with r-Net (Figure 6A). Comparative analysis revealed 266 differentially expressed genes (DEGs) (FDR <0.05; abs log FC >0.1) (Figure 6B), with upregulation of genes associated with naïve pluripotency and self-renewal (*Nanog*, *Esrrb*, *Klf4*), and downregulation of genes implicated in early mesodermal differentiation (*Dnmt3b*, *Gja1*) (Figure 6B)(Zanconato et al., 2015). A similar experimental approach was applied to human ESCs (hESCs), yielding consistent results: r-Net treatment in G1-phase hESCs regulated 1,566 genes (FDR <0.05; Abs Log FC >0.25) (Figure 6C). In line with our previous results, Gene Set Variation Analysis (GSVA) showed that r-Net tends to activate Yap1 score in both mouse and human G1-phase ESCs (Figure 6D). Furthermore, r-Net treatment reduced the cell cycle–dependent transcriptional heterogeneity of NNU target genes (Figure 6E). To complement these findings, we conducted a reciprocal loss-of-function experiment using siRNA-mediated knockdown of Netrin-1 in Fucci mESCs. Bulk, G1, and S–G2/M populations were sorted by FACS and subjected to RNA-seq (Figure 6F). Knockdown of Netrin-1 led to a specific downregulation of NNU target genes and Yap1-responsive transcripts in S–G2/M cells (Figure 6G-H). Collectively, these results demonstrate that NNU signaling shapes the transcriptome of both mouse and human ESCs in a cell cycle–dependent manner. Netrin-1 treatment in G1 is sufficient to restore the expression of key naïve pluripotency genes, suggesting that transient induction of NNU can be leveraged to modulate cell state and transcriptional identity in G1.

**Figure 6:**
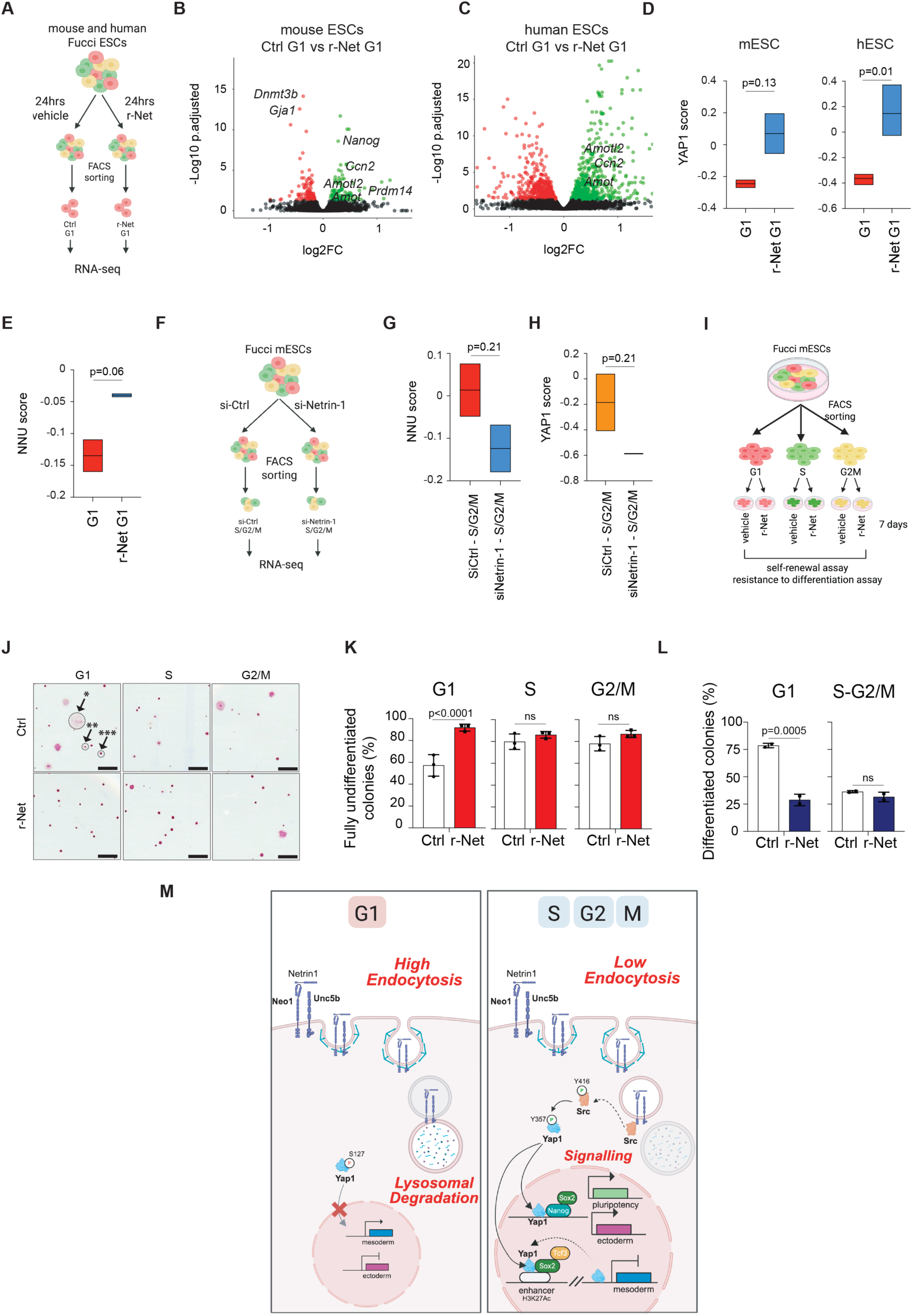
NNU signaling shapes the transcriptomes of mouse and human pluripotent stem cells to promote self-renewal of G1 ESCs. (**A**) Schematic representation of the experimental design for RNA-seq analysis on FACS sorted (1) G1, (2) S-G2/M and (3) G1 mESCs treated with r-Net. (**B**) Volcano plot showing differentially expressed genes between Ctrl G1 and r-Net G1 mouse ESCs. (**C**) Volcano plot showing differentially expressed genes between Ctrl G1 and r-Net G1 human ESCs. (**D**) GSVA Yap1 score in the indicated samples from mouse (left panel) and human (right panel) ESCs. Student T-test was used. (**E**) GSVA score for NNU-target genes (1552 genes significantly induced following Netrin-1 induction in mESC) in the indicated samples. Student T-test was used. (**F**) Scheme depicting the experimental design used to deplete Netrin-1 in S/G2/M phases of the cell cycle. (**G-H**) GSVA score calculation in the indicated samples. Student T-test was used. (**I**) Scheme depicting the experimental design used to restore Netrin-1 levels in G1. (**J**) Representative brightfield images of self-renewal assay. Cells were stained for Alkaline Phosphatase (AP). See methods for details. r-Net: recombinant Netrin-1. Scale bar: 800µm. On the top left picture, the *, ** and *** mark the position of a differentiated, mixed and fully undifferentiated colony, respectively. (**K**) Quantification of fully undifferentiated colonies from (**j**) in self-renewal assays. n= 3 independent experiments. Data are the mean +/- sd and two-way ANOVA-test was used for statistics. (**L**) Quantification of differentiated colonies in exit from pluripotency assays. n= 3 independent experiments. Data are the mean +/- sd and two-way ANOVA-test was used for statistics. (**M**) Model depicting the main findings of the study.

### Recombinant Netrin-1 Enhances Self-Renewal Capacity of G1-Phase ESCs

We next investigated whether recombinant Netrin-1 (r-Net) could functionally improve the self-renewal and differentiation resistance of mESCs in a cell cycle–dependent manner. Fucci mESCs were FACS-sorted into G1-, S-, and G2/M-phase populations and replated at low density in standard self-renewing conditions, in the presence or absence of r-Net (Figure 6I). Alkaline phosphatase (AP) staining, combined with quantification of undifferentiated and mixed colonies, revealed that G1-phase cells exhibited significantly reduced self-renewal capacity compared to their S- and G2/M-phase counterparts, consistent with previous reports (Figure 6J-K)(Pauklin and Vallier, 2013). Strikingly, r-Net treatment rescued the self-renewal potential of G1-phase ESCs, restoring their capacity to form fully undifferentiated colonies to levels comparable with S/G2/M-phase cells (Figure 6J-K). In contrast, r-Net treatment did not significantly affect the colony-forming ability of S- or G2/M-phase cells, consistent with the already elevated endogenous NNU activity in these phases. To assess differentiation resistance, FACS-sorted G1 and S/G2/M ESCs were replated in differentiation-inducing conditions (LIF withdrawal) for 7 days. As expected, G1-phase cells exited the pluripotent state more readily than S/G2/M cells. However, r-Net treatment rendered G1 cells as refractory to differentiation as cells in S/G2/M (Figure 6L). These findings demonstrate that the cyclic nature of NNU signaling contributes to the differential self-renewal and differentiation susceptibility of ESCs across the cell cycle. Moreover, transient supplementation with recombinant Netrin-1 is sufficient to enhance the functional properties of G1-phase ESCs, offering a strategy to stabilize naïve pluripotency across all cell cycle phases.

## Discussion

This study reveals that receptor internalization and endosomal trafficking represent a previously unrecognized, cell cycle–coupled mechanism for regulating stem cell fate and abscission dynamics. Using a CRISPR-engineered tri-color reporter ESC line, we track dynamic expression of the Netrin-1/Neo1/Unc5b (NNU) ligand–receptor complex at single-cell resolution. We find that NNU components display intrinsic heterogeneity that defines interconvertible stem cell subpopulations with distinct self-renewal capacity and lineage bias—NNU-high cells favor ectoderm/PGC fates, while NNU-low cells are mesoderm-biased and less self-renewing.

Strikingly, NNU protein levels oscillate throughout the cell cycle, peaking during S/G2/M and sharply declining in G1—not through changes in transcription or translation, but *via* tightly regulated clathrin-mediated endocytosis and lysosomal degradation. This periodic receptor turnover is conserved in mouse and human ESCs and is distinct from canonical pathways, positioning membrane trafficking as a new layer of pluripotency control. While Netrin-1 and its receptors have traditionally been characterized as signaling molecules, our findings indicate that they may also participate in broader cellular functions. These include the formation of protein complexes with biomechanical roles, notably in the context of cytokinetic abscission. It is plausible that components of the membrane complexes involved in abscission are internalized into late endosomes and asymmetrically inherited by daughter cells. Such asymmetry in endosomal content could confer a selective advantage to the daughter cell capable of more rapidly mobilizing these resources, potentially enhancing its responsiveness to environmental signals and influencing subsequent cell fate decisions.

Importantly, internalized NNU receptors activate a Src–Yap1 signaling axis from early endosomes, which reshapes Yap1 genomic binding and transcriptional outputs. This redistribution promotes naïve pluripotency and suppresses differentiation-associated genes in a cell cycle–specific manner. Perturbing receptor endocytosis delays cytokinetic abscission, further linking trafficking to fundamental cell division events. Finally, restoring Netrin-1 in G1-phase ESCs enhances self-renewal, and mimics the transcriptional state of S/G2/M cells.

Previous reports connected the cell cycle machinery and the control of pluripotency *via* a “nucleocentric model” in which cell cycle regulators, including the regulators cyclins and p53, control pluripotency features (Gonzales et al., 2015; Pauklin and Vallier, 2013). This model excluded endosomal trafficking and cyclicity in ligand/receptors activity to account for stem cell fate decisions (Dalton, 2015). We therefore propose a refined paradigm dynamically linking cell cycle, mechanics, endosomal trafficking and stem cell fate as depicted in Figure 6m. Together, these findings establish that dynamic receptor trafficking is not merely a means of signal attenuation but serves as a central mechanism for modulating stem cell identity in synchrony with the cell cycle that can be harnessed in a non-invasive manner for regenerative medicine.

## Acknowledgements

This work was supported by Fondation Bettencourt Schueller (Impulscience 2022 to FL), La Ligue Contre le Cancer Nationale et Régionale (to F.L.), Institut National du Cancer (2019-L22 to F.L.), Agence Nationale de la Recherche (Stemnet and Perionet to F.L, P.S and C.C), Fondation ARC (to A.H.), Centre Léon Bérard (F.L.), Labex DevWeCan (ANR-10-LABX-0061 to F.L and PS), Labex Cortex (ANR-11-LABX-0042 to P.S and F.L), LabEx REVIVE (ANR-10-LABX-0061 to P.S.), Institut Convergence PLAsCAN (ANR-17-CONV-0002) and Fondation pour la recherche médicale (N.C. and EQU202303016295 to P.S.) as well as the Deutsche Forschungsgemeinschaft (DFG:FOR2722 and SFB 1607, Z1 project for MK). We thank the core facilities of the Centre de Recherche en Cancérologie de Lyon (CRCL) and Centre Léon Bérard (CLB), and in particular Cyril Degletagne (Cancer Genomics Platform, CGP, CRCL) for single cell RNA-seq from Figure 1 and Thibault Andrieu (Flow Cytometry Core Facility, CYLE, CRCL) for FACS analysis related to Figure 4. We acknowledge Frédéric Delolme and Adeline Page from Protein Science Facility of SFR Biosciences (UAR3444/CNRS, US8/Inserm, ENS de Lyon, UCBL) for Mass Spectrometry analysis. We thank Sebastien Dussurgey from SFR Biosciences (Universite Claude Bernard Lyon 1, CNRS UAR3444, Inserm US8, ENS de Lyon) (PHENOCAN Equipex, ANR-11-EQPX-0035), for Figure 3. We thank C. Luccardini and D. Ressnikoff, CIQLE imaging center, SFR Santé Lyon-Est (UMS3453) for their help with image acquisition and analysis. We are grateful to Anne Vincent (CRCL) for her valuable assistance and expertise with the sucrose gradient technique. We thank Brigitte Manship for proofreading the manuscript.

## Author contributions

D.L performed most of the experiments presented in the figures. N.C. and X.G optimized and generated the TriC mESCs and conducted the initial sets of experiments related to Figures 1–3 and 5. E.Z. and H.H.V. performed the bioinformatics analyses. E.F. and A.H. conducted experiments related to Figures 4, 5 and 6. M.U, E.P and K.C provided cellular models for the study. C.R., I.A. and P.S. contributed to the experiments with the human Fucci ESCs and mitotic synchronisation. M.C. contributed to the generation of data for the study. N.G. and F.F. conducted the histologic analyses of teratoma. D.L, N.C, E.C, E.F and F.L designed experiments for the manuscript. D.L, E.C, E.F and F.L wrote the manuscript. F.L designed and supervised the study. All authors approved of and contributed to the final version of the manuscript.

---

**Figure S1:**
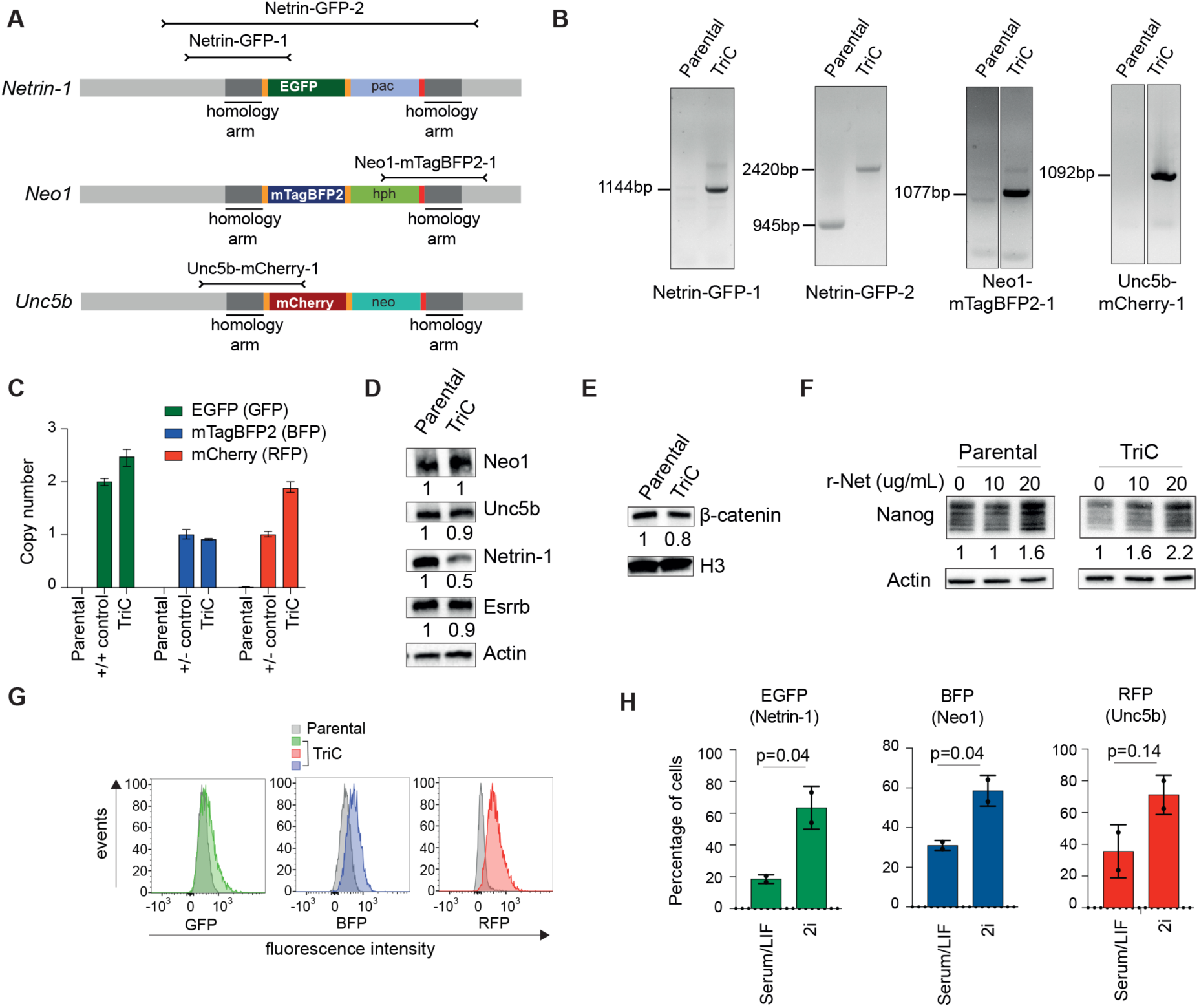
NNU level disciminate mESCs with various self-renewal and differentiation abilities. (**A**) Diagram of the Netrin-1, Neo1 and Unc5b loci after insertion of the reporter cassette by CRISPR/cas9 knock-in. The arrows indicate the amplicons generated by PCR on genomic DNA to evaluate the correct integration. Primer sequences are available in Supplementary Table 1. (**B**) Representative results of PCR conducted on genomic DNA of parental Cgr8 and TriC mESCs. Primers used are described in (A). (**C**) Copy number assays. PCR was conducted on genomic DNA to evaluate the number of copies of the transgenes integrated in the genome. See Methods for details. (**D-E**) Western blot for the indicated proteins in parental Cgr8 and TriC mESCs. (**F**) Western blot for Nanog. Parental Cgr8 and TriC mESCS were treated with recombinant Netrin-1 at the indicated times and concentrations(Huyghe et al., 2020). (**G**) Representative FACS plots of monofluorescence intensity in TriC mESCs with respect to the Parental Cgr8. Left: GFP – Netrin-1; Middle: Blue fluorescent protein BFP (mtagBFP2) – Neo1; Right: Red fluorescent protein RFP (mCherry) – Unc5b. One experiment representative of three independent experiments. (**H**) Graph depicts the percentage of cells in each culture conditions. Data are the mean +/- sd. n = 2 independent experiments. Student T-test was used.

**Figure S2:**
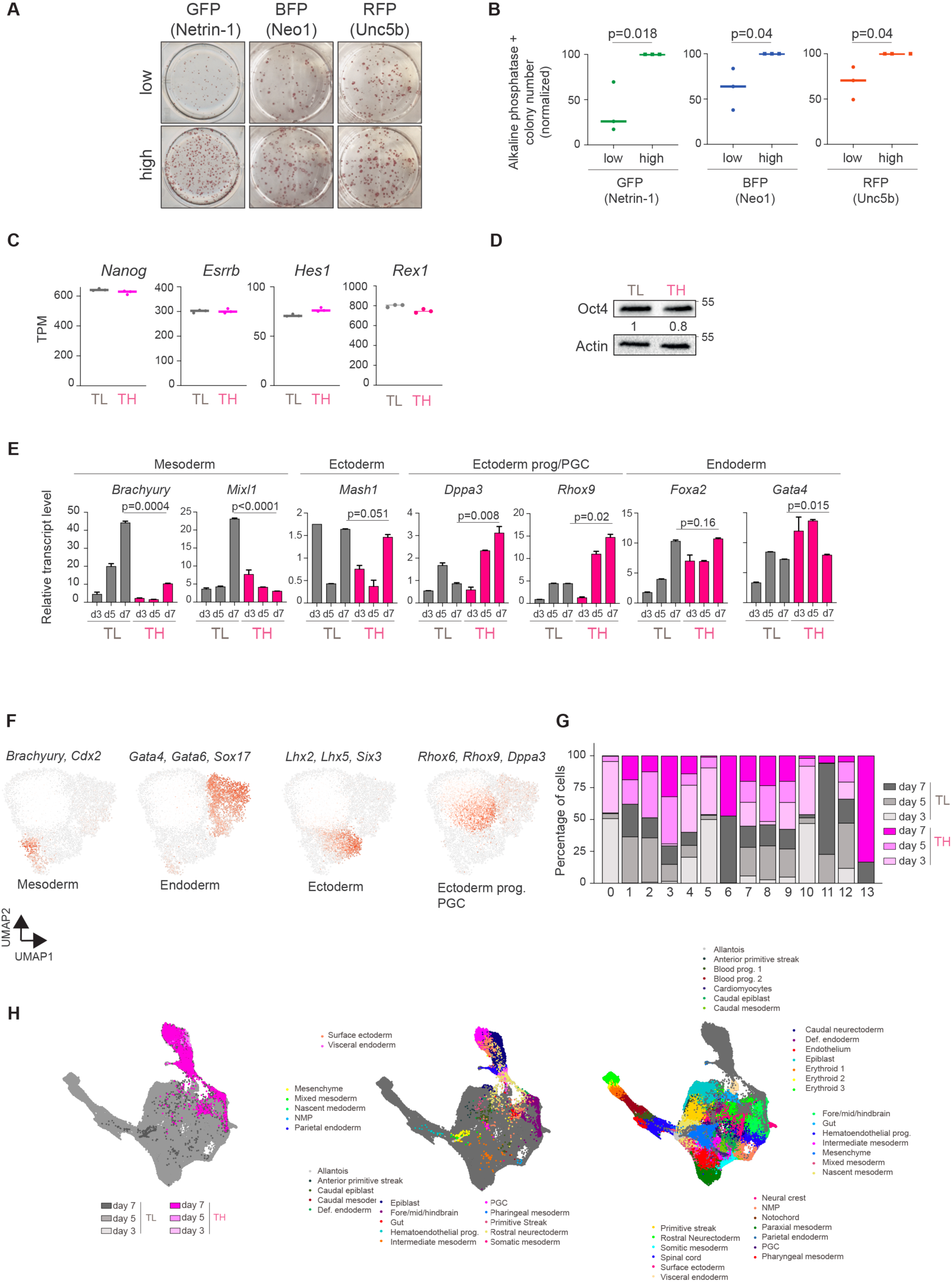
NNU level delineates mESCs with different epitranscriptomic features connected to cell cycle regulators. (**A**) Representative pictures of colony formation assay. TriC mESCs are FACS sorted on monocolor and replated at similar densities in self-renewing conditions. See methods for details. (**B**) Quantification of (a). n = 3 independent experiments. Data are the mean +/- sd and Student t-test was used for statistics. (**C**) Graph depicting *Nanog*, *Esrrb*, *Hes1* and *Rex1* transcript levels in FACS sorted NNU Triple High (TH) and Triple Low (TL). n= 3 independent experiments. Data are the mean +/- sd. TPM: Transcripts per Million. (**D**) Western blot for Oct4 in FACS-sorted TH and TL mESCs. (**E**) Bar charts showing transcript levels, as measured by RT-qPCR, during the differentiation of TH and TL cells. TH and TL cells were FACS sorted and subjected to embryoid body formation assay. n = 3 independent experiments. Data are the mean +/- sd and Student t-test was used for statistics. (**F**) Uniform Manifold Approximation and Projection (UMAP) visualization of scRNA-Seq profiles. TH and TL cells were FACS sorted and subjected to embryoid body formation. The graph integrates the replicate values of 20,849 pre-processed cells (individual dots), corresponding to three time points of differentiation processed in one sequencing experiment. The expression levels of the indicated marker genes are highlighted in red. (**G**) Percentage of cells in each of the 13 clusters defined in Figure 1. (**H**) Single-cell reference projection of TH and TL cells subjected to EB differentiation onto the atlas of gastrulating embryos (Pijuan-Sala et al., 2019), *de novo* visualization after merging the reference and query, an approach allowing for visualization of cell states that are present in the query dataset but are not represented in the reference.

**Figure S3:**
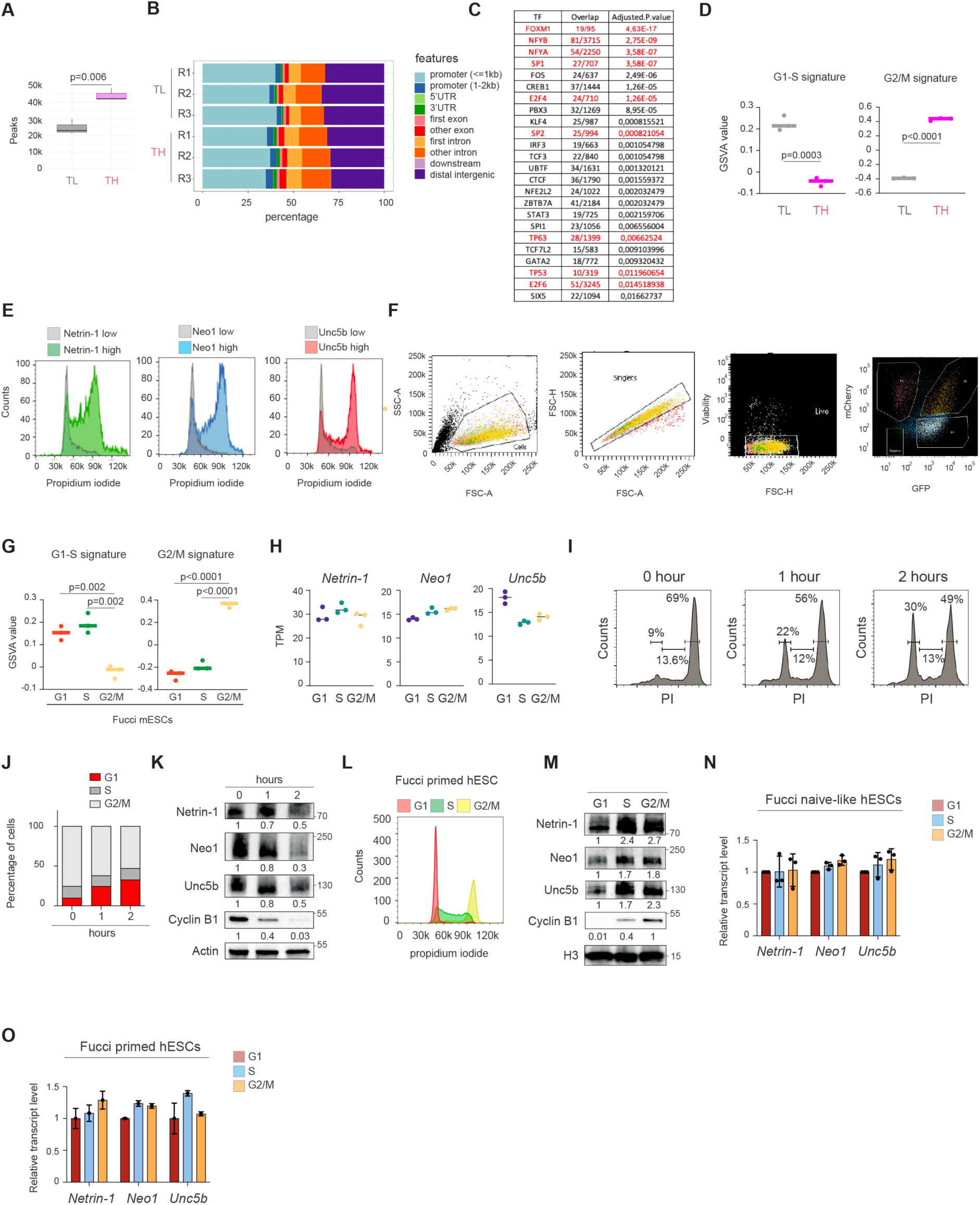
NNU levels discriminate mESCs with different epitranscriptomic features connected to cell cycle regulators. (**A**) Histogram of the number of ATAC peaks in TH and TL cells. (**B**) Distribution of the ATAC peaks on the genome. (**C**) Table depicting the prediction of master regulators of TH and TL cells. A list of 217 genes, upregulated and more accessible in TH cells, was extracted from Figure 2f and tested for ORA of ENCODE/ChEA consensus TF terms. (**D**) GSVA analysis of TH and TL cells. G1/S score was calculated using the FISCHER_G1_S_CELL_CYCLE geneset from the C2 collection of the Molecular Signature Database (Mootha et al., 2003; Subramanian et al., 2005)and G2/M score was calculated using the FISCHER_G2_M_CELL_CYCLE geneset from the aforementioned collection C2. Data are the mean +/- sd of 3 independent replicates and Student t-test was used for statistics. (**E**) Representative cell cycle profile of monocolor sorted TriC mESCs. Left panel: Netrin-1 GFP; Middle panel: Neo1-BFP; right panel: Unc5b-RFP. Cell cycle analysis was conducted with propidium iodide. (**F**) FACS gating strategy for Fucci mESCs. (**G**) GSVA analysis of Fucci mESCs. Data are the mean +/- sd of 3 independent replicates and Student t-test was used for statistics. (**H**) *Netrin-1*, *Neo1* and *Unc5b* transcript levels in Fucci mESCs FACS sorted in G1, S or G2/M. Transcriptomic data were used. n = 3 independent experiments. (**I**) Representative cell cycle profile of mESCs during the release of the mitotic block induced with Demecolcine. See methods for details. (**J**) Graph depicting the percentage of mESC in each phase of the cell cycle in same settings as (**I**). Data correspond to one experiment representative of 2 independent experiments. (**K**) Western blot for the indicated proteins in similar settings as (I-J). (**L**) Representative cell cycle profile of Fucci primed hESC. Cells were FACS sorted in G1, S and G2/M and analysed for cell cycle using propidium iodide to demonstrate the efficacy of the Fucci system. (**M**) Western blot for the indicated proteins. (**N-O**) *Netrin-1*, *Neo1* and *Unc5b* transcript levels in Fucci hESCs FACS sorted in G1, S or G2/M. RT-qPCR data are presented. (**N**) Fucci naïve-like hESCs. Data are the mean +/- sd. n = 3 independent experiments. (**O**) Fucci primed hESCs. Data are the mean +/- sd. n = 2 technical replicates.

**Figure S4.**
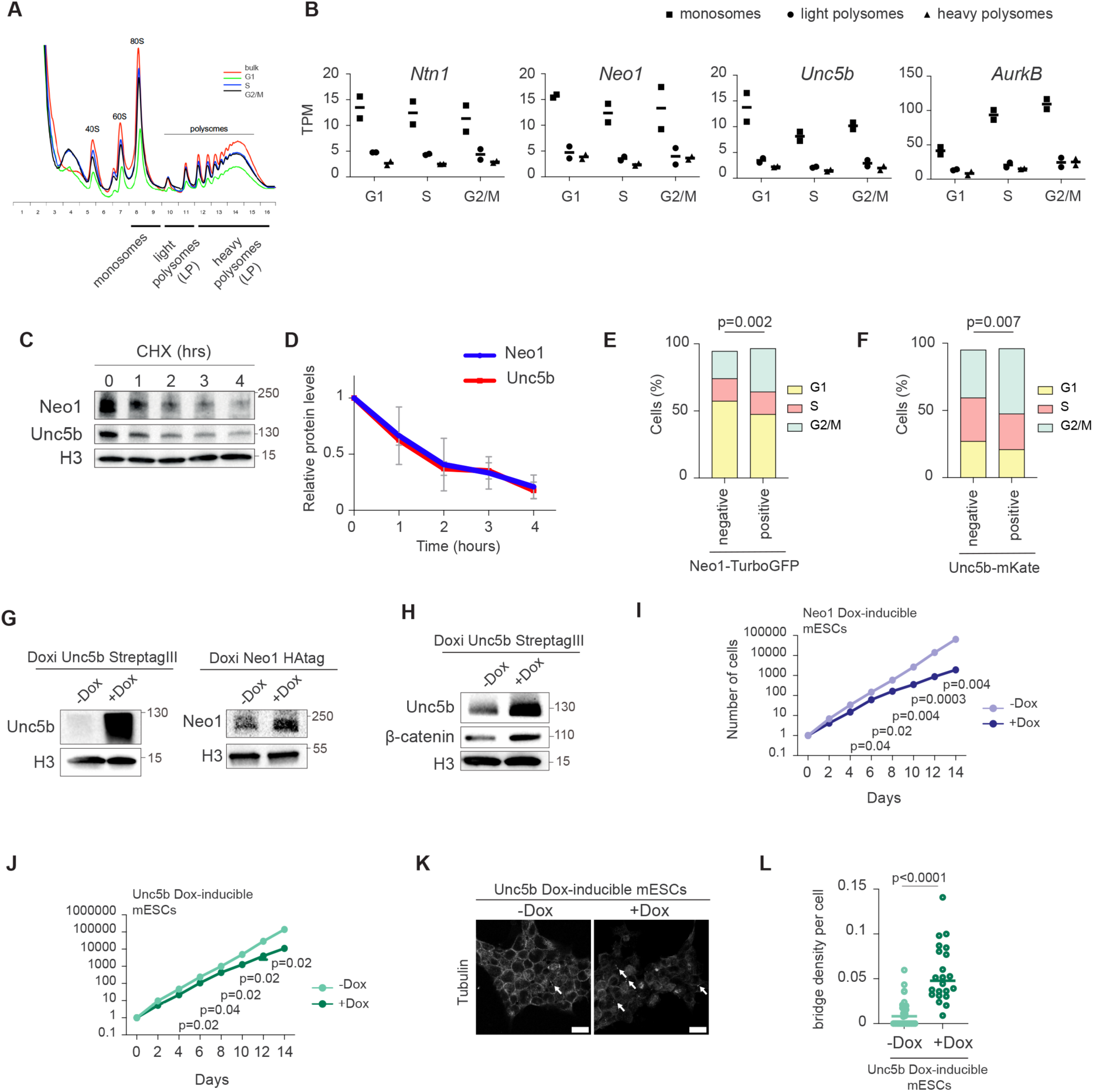
(**A**) Representative example of polysome isolation by gradient centrifugation conducted on bulk and FACS-sorted mESCs in G1, S and G2/M. (**B**) Loading of *Ntn1*, *Neo1*, *Unc5b* and *AurkB* transcripts on heavy and light polysomes during the cell cycle. Graph depicts transcript levels (RPKM) in heavy and light polysomes fractions isolated from samples described in (**A**). (**C**) Western blot for Neo1 and Unc5b in mESCs treated or not with Cycloheximide (CHX) for the indicated time. (**D**) Quantification of Neo1 and Unc5b protein stability. Graph depicts data from (**c**). n= 3 independent experiments. (**E-F**) Graph depicting the percentage of cells in each phase of the cell cycle in mESCs. (E) mESCs expressing or not the Neo1-TurboGFP construct. (F) mESCs expressing or not the Unc5b-mKate construct. Data correspond to 3 independent experiments. (**G**) Western Blot for Unc5b (left panel) and Neo1 (right panel) in Dox-inducible mESCs. (**H**) Western Blot for Unc5b and β-catenin in Unc5b-StreptagIII Dox-inducible mESCs in non-treated and doxycycline-treated condition. (**I**) proliferation curve of Neo1 Dox-inducible mESCs for 14 days. (**J**) Proliferation curve of Unc5b Dox-inducible mESCs for 14 days. (**K**) Representative confocal image of mESCs expressing exogenously Unc5b stained for α-tubulin (white). Left panel: non treated Right panel: doxycycline treatment for 14 days. A maximum Z projection is shown. Scale bar: 20 µm. (**L**) Dot plot showing the fraction of cells with tubulin bridges (number of bridges divided by number of cells in a given analysis frame) in Unc5b Dox-inducible mESCs non treated or treated with doxycycline for 14 days. Mean is shown. n = 3 independent experiments. Student T-test was used.

**Figure S5.**
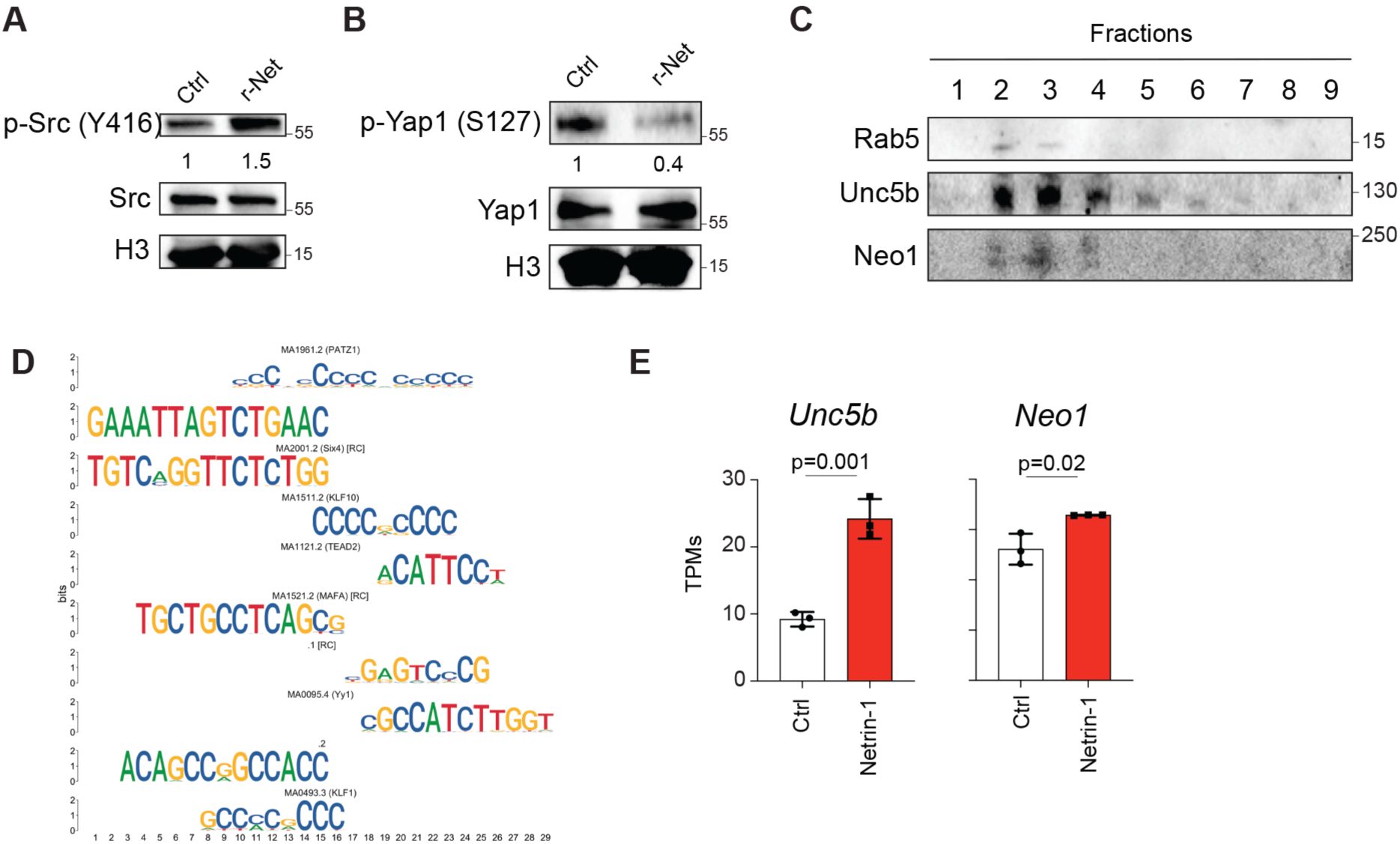
(**A**) Western Blot for phosphorylated Src on Tyr416 in mESCs treated or not with recombinant Netrin-1. (**B**) Western Blot for phosphorylated Yap1 on Ser127 in mESCs treated or not with recombinant Netrin-1. (**C**) Western Blot of cell fractions obtained by sucrose gradient fractionation. (**D**) MEME motif analysis from Yap1 ChIP-seq analysis. **(E)** Unc5b and Neo1 transcript levels in G1 Fucci mESC treated or not with recombinant Netrin-1.

## Methods

### Cell culture

mESCs Cgr8 ESC (ECACC 07032901) and E14-Fucci ESCs (kind gift from Savatier lab) were grown and stably transfected as described previously(Huyghe et al., 2020). For TriC ESCs generation, CRISPR knock-in was conducted following F. Zhang’s laboratory protocol(Ran et al., 2013). Briefly, cells were transfected with the pSpCas9(BB)-2A-puro plasmid (Addgene #48139) and the homology repair plasmid construct using Lipofectamin 2000 (Invitrogen). Selection was applied the day after transfection, and monoclonal colonies were selected and genotyped by genomic PCR before validation by copy number assay (TaqMan assay, Life technologies) and by cell sorting followed by Western blot. Netrin-1 exogenous expression was induced in Ntn1 inducible mESCs(Huyghe et al., 2020) by treating cells 24h with 1 µg/mL of doxycycline.

Endocytosis was inhibited by treating the cells with 0.5µg/mL Filipin III (Sigma), 15µM Chloroquine (Sigma) or 5µM Pitstop2 (Merck) for 24h. RhoA and Arp2/3 were inhibited treating the cells with 10 µM Rhosin hydrochloride and 50 µM CK666 respectively for the indicated time points. RhoA was induced in RhoA inducible mESCs by treating the cells with doxycycline at 1 µg/mL for 48h, as previously described(De Belly et al., 2021).

The PB-PuroR-CAG-mVenus:hGeminin-IRES-mCherry:hCdt1 plasmid was used to generate-FUCCI hESCs reporter cells, as described recently(Aksoy et al., 2021). Human Oscar ESCs were transfected using the NEON transfection system according the manufacturer’s instructions. Briefly, cells were dissociated and resuspended at 10.10^6^ cells/mL. For transfection, 100μL of the cell suspension was mixed with 2.5μg of the PB-PuroR-CAG-mVenus:hGeminin-IRES-mCherry:hCdt1 vector and 2.5 μg of PBase plasmid. Transfection parameters were 1,050 V, 20 ms, and 2 pulses. Cells were plated on growth-inactivated DR4 murine embryonic fibroblasts in medium supplemented with 10 μM ROCK inhibitor (Y-27632; Miltenyi Biotec) and selected in 1 μg/mL Puromycin.

Primed to PXGL conversion was performed using the protocol described(Bredenkamp et al., 2019). Briefly, primed cells were dissociated and plated on fresh feeder cells for 24 h before shifting to the PXGL culture media composed of N2B27 (in house) supplemented with 1 μM PD0325901, 2μM XAV939, 2 μM Gö6983 and 10 ng/mL human LIF. Both mouse and human ESCs were treated with r-Net as described previously(Huyghe et al., 2020).

For gene silencing, cells were transfected with 20nM of ON-TARGET plus siRNA (Horizon) and Lipofectamine 2000 following the manufacturer’s instructions and collected after 48-72 hours for protein and RNA extraction. siRNA used in this study are listed in Supplementary Table 1.

### Interconvertibility assay

For interconvertibility analysis, 250,000 TriC mESC sorted for Ntn1-High/Ntn1-Low; Neo1-High/Neo1-Low; Unc5b-High/Unc5b-Low were plated in MW6 coated with 0.1% of gelatin. For the first time point (0h) the cells were analyzed at the FACs just after sorting. For subsequent time points (24h and 48h), the cells were trypsinized before FACs analysis.

### Colony formation assay

For colony formation assay, ∼500 cells were plated in MW6 coated with gelatin 0.1% in the indicated medium conditions, and medium was changed every day for 7 days. The alkaline phosphatase-positive colonies were detected using the Alkaline Phosphatase Diethanolamine Activity Kit (AP0100-1KT, Sigma) and images were taken using the Epson Perfection V600 photo scanner.

### Differentiation assays

Differentiation into RSCs and EpiLCs was induced(Neagu et al., 2020) by plating 250,000 cells in MW6 coated with gelatin 0.1%. The medium was changed the day after with RSC differentiation medium (N2B27 medium supplemented with 2 µM IWP2 (681671, Sigma) and 1µM PD0325901 (444968, Sigma) or EpiLC differentiation medium (N2B27 medium supplemented with 20 ng/mL ActivinA (338-AC-010/CF, R&D Systems) and 12ng/mL bFGF (3139-FB-025/CF, R&D Systems)) and cells were collected 48h after for subsequent analysis. For EBs differentiation, TriC mESCs were FACs sorted for TH and TL cells and 280,000 cells were plated in Ultra-Low Attachment MW6 (CLS3471-24EA, Sigma) in DMEM supplemented with 7.5% FBS, 4 mM L-Glutamine, 100 units/mL Penicillin-Streptomycin and samples were collected at indicated time points.

### Protein extraction, Western blot and protein stability assays

Cells were collected and lysed in RIPA buffer (50mM Tris-HCl pH 8.0, 150mM NaCl, 1% NP40, 0.5% Sodium Deoxycholate, 0.1% SDS) supplemented with Phosphatase Inhibitor Cocktail 2/3 (Sigma) and cOmplete™ Protease Inhibitor (Sigma) for 30 min on ice. Cell debris were removed by centrifugation for 10 minutes at 10,000 g at 4°C and protein quantification was performed with the BCA Protein Assay Kit (Life Technologies). Actin or Histone H3 were used as loading control. The antibodies used for proteins detection are listed in Supplementary Table 1. For protein stability analysis, 700,000 cells were plated in MW6 coated with 0.1% of gelatin and treated with 5µg/mL Cycloheximide (CHX). Cells were collected by scraping at the indicated time points and proteins were extracted in RIPA buffer.

### RNA isolation and RT–qPCR

RNA was extracted using Trizol (Sigma) following manufacturer instructions. cDNA was generated using the RevertAid H Minus First Strand cDNA Synthesis Kit (Life Technologies) and RT-qPCR reactions were performed using the Syber Green master mix (Roche). Results were normalized to the housekeeping gene Actin. Primers used are listed in Supplementary Table 1.

### FACS analysis and sorting

Flow cytometry was carried out on a LSRFortessa flow cytometer and analyzed by FlowJo software. For Cell-cycle distribution analysis, cells were fixed with 70% ethanol overnight before incubation with 500ng/mL Propidium Iodide (Sigma) for 15 minutes at room temperature. Cell sorting was performed at 4°C with the BD FACS Aurora sorter. After sorting, the cells were centrifugated at 4°C before being plated in fresh medium and collected at different time points or used for protein/RNA extraction, cell cycle analysis and interconvertibility assays.

### Endocytosis assay

1×106 cells were seeded in MW6 24h before treatment. Cells were first incubated 30 min with 6µg of Alexa Fluor Dextran (Dextran, Alexa Fluor 647; 10,000 MW, Thermofischer Scientific). Cells were then rinsed and trypsinated before FACS analysis (Fortessa Horizon). We also performed a cytometry analysis coupled with image capture using ImageStreamX Mk II imaging cytometer Cytek. The analysis was carried out with IDEAS software version 6.2. The 40x magnification was used to selectively visualize cells in focus. We used logarithm bright detail intensity to analyze dextran-AF647 positive cells in G1, S and G2/M phases.

### Time lapse

A total of 200,000 Neo1–Turbo-GFP or Unc5b–mKate cells were seeded onto 35 mm ibidi dishes precoated with poly-L-ornithine (Merck) 48 h prior to imaging. Time-lapse experiments were performed using the Nanolive 3D Explorer system, with fluorescence images excitation at 488 nm (Turbo-GFP) or 555 nm (mKate) together with bright-field images, every 10 min over a 24 h period.

### Protein stability

For protein stability analysis, 700,000 Control KO and Netrin KO cell lines were plated in MW6 coated with 0.1% gelatin and treated with 5μg/mL cycloheximide and cells were collected at different time points. To determine whether receptor degradation was related to lysosome degradation, cells were treated 24 h prior to cycloheximide treatment with 15uM of chloroquine. We performed western blot to evaluate Neogenin 1 and Unc5b expression stability in presence or in absence of ligand.

### α-tagged Netrin-1 stable cell line

mESCs Cgr8 ESC were grown and stably transfected as described previously 6. Cells were stably tranfected with KV4552 Full netrin plasmid from Manuel Koch’s lab using Lipofectamin 2000 (Invitrogen). Polyclonal cells were selected with puromycin at 1ug/ml during 72h. α-tagged Netrin-1 exogenous expression was induced by treating cells 24h with 1 μg/mL of doxycycline.

### Time lapse to follow the degradation of α-tagged Netrin-1 by lysosome

α-tagged Netrin-1 cells were plated at 0.1M cells 48h before the start of the film in 35mm ibidi dishes coated with poly-L-ornithine (Merck) and cultured in medium containing 1ug/ml doxycycline. 1h before the start of the film, we removed the medium and incubated the cells for 1h on ice with the antibody solution containing FluoTag-X2 anti-ALPHA single domain antibody coupled with AF647 diluted 1:500 (Abnova ref RAB00922) and lysotracker ex/em 504/511 diluted 1:1000 (life technology ref L7526) in cold PBS1X. Just before starting the film, remove the antibody solution and add ES medium at 37°C without red phenol. Take a photo every 2 minutes for 30mins Pictures acquisition were performed with Andor Spinning Disk and merged on FIJI imageJ software. All quantification were performed with QuPath software (Bankhead, P. et al. QuPath: Open source software for digital pathology image analysis. Scientific Reports (2017). https://doi.org/10.1038/s41598-017-17204-5). Lyso-Tracker detection was performed with the Water Shed Cell Detection tool, α-tagged Netrin-1 with the Positive Cell Detection and Endocytosis was quantify by using an Object Classifier JSON.

### Endosome isolation

The protocol is based on ref.(Walker et al., 2016). mESCs were plated in MW6 pre-coated with 0.2% gelatin. The following day, at approximately 80% confluency, cells were treated with 10µg/ml recombinant Netrin for 30 minutes. After treatment, cells were washed with PBS and harvested by centrifugation at 1000rpm for 5 minutes at 4°C. The cell pellets were resuspended in 500µl of buffer 1 (8.5% of sucrose, 1mM EDTA), and gently dissociated 10X with a 23G seringe. Nuclei were removed by centrifugation at 1,000 g for 5 minutes, and the supernatant fraction was collected, to which 500µl of buffer 2 was added (41.5% of sucrose, 1mM of EDTA). 25–45% sucrose gradients were poured using the Gradient Master (Serlabo Technologies). After ultracentrifugation at 100 000 g during 1hr at 4°C on a SW41 Beckman rotor, fractions of 700µl each were collected from each gradient using an ISCO UA-6 detector.

### Cell synchronization

For synchronization in G1 phase, cells were grown in N2B27+2i+LIF(Smith, 2017) treated with 40µM of LY294002 PI3K inhibitor as previously described(Jirmanova et al., 2002) and collected after 24h for subsequent analysis. To induce mitotic block, cells were treated according to previous published protocol(Savatier et al., 2002).

### RNA-seq sample processing and analysis

For Bulk RNA-seq, samples were processed with Macherey-Nagel™ ARN NucleoSpin™ kit and RNAs were extracted following manufacturer instructions. RNA samples were quantified using Qubit 4.0 Fluorometer (Life Technologies, Carlsbad, CA, USA) and RNA integrity was checked with RNA Kit on Agilent 5300 Fragment Analyzer (Agilent Technologies, Palo Alto, CA, USA).

RNA sequencing libraries were prepared using the NEBNext Ultra RNA Library Prep Kit for Illumina following manufacturer’s instructions (NEB, Ipswich, MA, USA). Briefly, mRNAs were first enriched with Oligo(dT) beads. Enriched mRNAs were fragmented for 15 minutes at 94 °C. First strand and second strand cDNAs were subsequently synthesized. cDNA fragments were end repaired and adenylated at 3’ends, and universal adapters were ligated to cDNA fragments, followed by index addition and library enrichment by limited-cycle PCR. Sequencing libraries were validated using NGS Kit on the Agilent 5300 Fragment Analyzer, and quantified using Qubit 4.0 Fluorometer (Invitrogen, Carlsbad, CA).

The sequencing libraries were multiplexed and loaded on the flowcell on the Illumina NovaSeq 6000 instrument according to the manufacturer’s instructions. The samples were sequenced using a 2×150 Pair-End (PE) configuration v1.5. Image analysis and base calling were conducted by the NovaSeq Control Software (NCS). Raw sequence data (.bcl files) generated from Illumina NovaSeq was converted into FASTQ files and de-multiplexed using Illumina bcl2fastq program version 2.20. One mismatch was allowed for index sequence identification. The FASTQ files were adapter-trimmed using the bbduk.sh procedure from the BBMap suite using its default reference set of adapters, and quasi-mapped to the ENSEMBL’s(Martin et al., 2023) GRCm39 (release 109) in the case of mouse and ENSEMBL’s GRCh38 (release 109) in the case of human transcriptomes using salmon v1.10.1(Patro et al., 2017). The salmon index in both cases was generated with full sets of decoys (“SAF”) to confer the highest mapping accuracy possible.

### GSVA analyses

GSVA analyses were performed using the GSVA package in R(Hanzelmann et al., 2013) and since the GSVA scores are distributed quasi-normally, we applied the t-test of statistical significance to them.

### ATAC-seq sample processing and analysis

Cells were harvested and frozen in culture media containing FBS and 5% DMSO. Cryopreserved cells were sent to Active Motif to perform the ATAC-seq assay. The cells were then thawed in a 37°C water bath, pelleted, washed with cold PBS, and tagmented as previously described(Buenrostro et al., 2013), with some modifications(Corces et al., 2017). Briefly, cell pellets were resuspended in lysis buffer, pelleted, and tagmented using the enzyme and buffer provided in the Nextera Library Prep Kit (Illumina). Tagmented DNA was then purified using the MinElute PCR purification kit (Qiagen), amplified with 10 cycles of PCR, and purified using Agencourt AMPure SPRI beads (Beckman Coulter). Resulting material was quantified using the KAPA Library Quantification Kit for Illumina platforms, and sequenced with PE42 sequencing on the NovaSeq 6000 sequencer.

The NGS data are deposited under the GEO accession number GSE267978 (private token for reviewers ibujgckwhdutpaz).

### SC-RNA-seq sample processing and analysis

EBs were collected at different time points and dissociated with TrypLE Express Enzyme (Life Technologies) and samples were processed for sc-RNA seq as previously described(Huyghe et al., 2022).

#### Raw Data

Raw data from Cellranger yielded six samples, each containing varying numbers of cells ranging from 8,098 to 24,457. To ensure data integrity and reliability, a rigorous quality control process was implemented. Initially, we employed visual inspection through violin plots to filter out potentially problematic cells. Criteria for exclusion included cells with 200,000 transcripts or more, those with mitochondrial transcripts exceeding 10%, and those with fewer than 500 unique genes. Moreover, a distinct bimodal distribution emerged in both the number of transcripts and detected genes per cell. The lower mode in these distributions was indicative of dying or ruptured cells. To enhance the robustness of subsequent analyses, only cells constituting the upper mode were retained, defining a threshold of at least 15,000 transcripts per cell.

#### Downsampling and Outlier Removal

Post-filtering, the retained cells ranged from 49% to 61% of the initial count per sample, resulting in a final cell count between 3,959 and 12,382 cells. To standardize the dataset for subsequent analyses and ensure comparability, all samples were downsampled to 3,500 cells, aggregating a total of 21,000 cells across all samples. Further analysis of the Uniform Manifold Approximation and Projection (UMAP) plot for these 21,000 cells revealed 141 outlying cells, disconnected from any identifiable cluster. To preserve the integrity of the overall embedding and prevent potential distortions induced by these outliers, they were systematically removed. Consequently, the dataset was refined to 20,859 cells distributed across the six samples. All analyses were performed using Seurat v4(Hao et al., 2021) in R v4.3.1.

#### Data Homogeneity and Sample Retention

Following the removal of outliers, each sample retained between 98.0% and 99.9% of its initial cells. Given this negligible loss and the impracticality of re-equalizing cell counts between samples, it was deemed unnecessary to pursue further adjustments. This stringent quality control pipeline ensures the reliability and consistency of the dataset, laying a solid foundation for subsequent analyses and interpretations.

### Immunoprecipitation followed by mass spectrometry

#### Immunoprecipitation

mESCs were collected and lysed in cool IP lysis Buffer (20mM Tris pH7.5, 150mM NaCl, 1% NP40, proteases /phosphatases inhibitors). The immunoprecipitation assays were conducted using at least 1mg of protein lysate incubated with magnetic beads directed against Strep-tag III (IBA MagStrep XT beads) rotating overnight at 4°C. The next day, the immunocomplexes were washed three times with distilled water using a magnetic rack. Proteins were eluted in 2x Loading Buffer by boiling for 10 minutes. Samples were subsequently analyzed by Western Blot.

#### In-Solution enzymatic digestion

50 µL of IP samples were mixed with 50 μL of Lyse buffer provided in the PreOmics iST Kit (PreOmics GmbH; Martinsried, Germany). The samples were reduced and alkylated by incubation at 95 °C for 10 min and continuously shaken at 1000 rpm. Then samples were digested for 2h30min at 37°C and processed following the manufacturer’s protocol.

#### nanoLC-MS/MS analysis

The samples were analyzed using an Ultimate 3000 nano-RSLC (Thermo Scientific, San Jose California) coupled on line with a Q Exactive HF mass spectrometer via a nano-electrospray ionization source (Thermo Scientific, San Jose California). 1 µL of the peptide mixtures were loaded on a PepMap NEO C18 trap-column (300 µm ID x 5 mm, 5 µm, Thermo Fisher Scientific) for 3.0 minutes at 20 µL/min with 2% ACN, 0.05% TFA in H2O and then separated on a C18 Acclaim Pepmap100 nano-column, 50 cm x 75 μm i.d, 2 μm, 100 Å (Thermo Scientific) with a 60 minutes linear gradient from 3.2% to 40% buffer B (A: 0.1% FA in H2O, B: 0.1% FA in ACN), from 40 to 90% of B in 2 min, hold for 10 min and returned to the initial conditions in 1 min for 14 min. The total duration was set to 90 minutes at a flow rate of 300 nL/min. The oven temperature was kept constant at 40°C. Samples were analysed with a TOP20 HCD method: MS data were acquired in a data dependent strategy selecting the fragmentation events based on the 20 most abundant precursor ions in the survey scan (350-1600 Th). The resolution of the survey scan was 60,000 at m/z 200 Th. The Ion Target Value for the survey scans in the Orbitrap and the MS2 mode were set to 3E6 and 1E5 respectively and the maximum injection time was set to 60 ms for both scan modes. HCD MS/MS spectra acquisition parameters were as follows: collision energy = 27; isolation width of 2 m/z; precursors with unknown charge state or a charge state of 1 were excluded. Peptides selected for MS/MS acquisition were then placed in an exclusion list for 20s using the dynamic exclusion mode to limit duplicate spectra. The spray voltage was set in positive ion at 1800 V and ion transfer tube maintained at a temperature of 250°C.

#### Data Analysis

Proteins were identified by database searching using Sequest HT with Proteome Discoverer 2.5 software (Thermo Scientific) against Mus musculus uniprot database (2023-01 release, 102924 sequences) and a contaminant database. Precursor and fragment mass tolerance were set at 10 ppm and 0.02 Da respectively, and up to 2 missed cleavages were allowed. Oxidation (M), acetylation (Protein N-terminus), were set as variable modification, and Carbamidomethylation (C) as fixed modification. Full Trypsin was selected as digestion enzyme parameter. Peptides and proteins validation were performed by Percolator and a false discovery rate of 1% for peptides and proteins was set.

### Statistics and reproducibility

Western blots were quantified using ImageJ software. Statistical analyses of mean and variance were performed with GraphPad Prism and Student’s t-test or Anova tests where indicated. For western blots presented in the figures, three independent experiments gave similar results.

### Teratoma

TriC mESCs were FACS sorted prior to injection. Teratoma formation was conducted as previously described(Huyghe et al., 2020; Huyghe et al., 2022; Ozmadenci et al., 2015). For histological examination, tissue samples were fixed in 10% buffered formalin and embedded in paraffin. 4-µm-thick tissue sections of formalin-fixed, paraffin-embedded tissue were prepared according to conventional procedures. Sections were then stained with hematoxylin and eosin and were scanned with panoramic scan II (3D Histech, Hungary) at 20× magnification. Captured images were processed using HALO software (Indica Labs, V3.6.4134.345) and HALO AI (Indica Labs, V3.6.4134) for tissue classification. The MiniNet Deep Learning model was trained to segment each tissue type. Representative areas on the slides were annotated to train the model.

### Animal study

The animals were maintained in a specific pathogen-free animal facility P-PAC (small animal platform). All of the experiments were performed in accordance with the animal care guidelines of the European Union and were validated by the local Animal Ethics Evaluation Committee (C2EA-15 agreed by the French Ministry of High School and Research). Embryos were flushed and imaged as previously described(Huyghe et al., 2020).

