## Supplemental Table 1 for "A ligand/receptor trafficking clock governs self-renewal and abscission dynamics in pluripotent stem cells"

### Supplementary Table 1

| Antibody | Company | Reference | Dilution |
|---|---|---|---|
| Netrin 1 | Abcam | ab126729 | 1:1,000 |
| Neogenin 1 | Ozyme | 39447 | 1:1,000 |
| Unc5b | Ozyme | 13851 | 1:1,000 |
| β-actin–peroxydase | Sigma | A3854 | 1:10,000 |
| Anticorps p53 (Pab 1801) | Santa Cruz | sc-98 | 1:1,000 |
| Histone H3 | Abcam | ab1791 | 1:1,000 |
| Aurora-A Kinase | BD biosciences | 610938 | 1:1,000 |
| Bmp4 | Abcam | ab39973 | 1:1,000 |
| Fgf4 | Abcam | ab106355 | 1:1,000 |
| Fgfr1 | Ozyme | 9740T | 1:1,000 |
| GP130 | Ozyme | 3732S | 1:1,000 |
| β-catenin | Santa Cruz | sc-7963 | 1:1,000 |
| Cyclin B1 | Santa Cruz | sc-245 | 1:1,000 |
| Oct-3/4 | Santa Cruz | sc-5279 | 1:1,000 |
| Phospho-YAP1-S127 | Sigma | SAB5700367 | 1:1,000 |
| YAP1 | Santa Cruz | sc-101199 | 1:1,000 |
| LATS1 (C66B5) | Cell Signaling | 3477 | 1:1,000 |
| Nanog | Cell Signaling | 3580 | 1:1,000 |
| Esrrb | R&D system | PP-H6705-00 | 1:1,000 |

**siRNA**

| Name | Reference | Company |
|---|---|---|
| Non-targeting Pool siRNA | D-001810-10-20 | Horizon |
| Cltc siRNA | L-063954-00-0005 | Horizon |
| RhoA siRNA | L-042634-00-0005 | Horizon |
| Arpc2 siRNA | L-043464-01-0005 | Horizon |
| Arpc3 siRNA | L-040765-01-0005 | Horizon |

**oligo**

| Name | Sequence | Application | Species |
|---|---|---|---|
| Ntn1_F | CCCTTGCATCAAGATTCCTGTG | rt-qPCR | Mouse |
| Ntn1_R | CTTCATGTTCATCTTCAGCTTGCC | rt-qPCR | Mouse |
| Neo1_F | CCATCTAGCACCAGCCACAA | rt-qPCR | Mouse |
| Neo1_R | TTCTCAACACGCTCCCTAGC | rt-qPCR | Mouse |
| Unc5b_F | AACTACTGGCCAAGTACCAGGAGA | rt-qPCR | Mouse |
| Unc5b_R | GACCTTACAGGTGAACTCCGTG | rt-qPCR | Mouse |
| Brachyury_F | GAATTCGTCCACCCCCTGTC | rt-qPCR | Mouse |
| Brachyury_R | TAGGCATGTTCCAAGGGCAG | rt-qPCR | Mouse |
| Mixl1_F | AGTTGCTGGAGCTCGTCTTC | rt-qPCR | Mouse |
| Mlixl1_R | GTTCTGGAACCACACCTGGA | rt-qPCR | Mouse |
| Mash1_F | TTTGGAAGCAGGATGGCAGC | rt-qPCR | Mouse |
| Mash1_R | GGGCTTAGGTTCAGACACCA | rt-qPCR | Mouse |
| Dppa3_F | AGAGAAGGGTCCGCACTTTG | rt-qPCR | Mouse |
| Dppa3_R | AATGGCTCACTGTCCCGTTC | rt-qPCR | Mouse |
| Rhox9_F | CTTCCCCAGCTTGCGAGTAA | rt-qPCR | Mouse |
| Rhox9_R | CTGTAATCGGTGGCAGTTCG | rt-qPCR | Mouse |
| FoxA2_F | GGTCGTTTGTTGTGGCTGTT | rt-qPCR | Mouse |
| FoxA2_R | CTCACGGAAGAGTAGCCCTCG | rt-qPCR | Mouse |
| Gata4_F | TGGAAGACACCCCAATCTCG | rt-qPCR | Mouse |
| Gata4_R | TAGTCTGGCAGTTGGCACAG | rt-qPCR | Mouse |
| Ntn1_F | TGACGGGTATCACCTGCAAC | rt-qPCR | Human |
| Ntn1_R | AGGAATCGCAGTCTTCAGGC | rt-qPCR | Human |
| Unc5b_F | CAGGGCAAGTTCTACGAGAT | rt-qPCR | Human |
| Unc5b_R | TGGTCCAGCAGGATGTGA | rt-qPCR | Human |
| Neo1_F | ATGGTGACCAAAGGTCGAAG | rt-qPCR | Human |
| Neo1_R | AGTCACATCCTTGGGTGGAG | rt-qPCR | Human |
| Ntn1 | TGCAAGAAGGCCTAGCGCGG | CRISPR knock-in | Mouse |
| Neo1 | TTGGAGTCATGTCCAGGCGA | CRISPR knock-in | Mouse |
| Unc5b | CACTCAGCAATCGCCATCTG | CRISPR knock-in | Mouse |

List of NNU interactants

| Protein FDR Confidence: Combined | Master | Accession | Description | Contaminant | Marked as | Sum PEP Score | Coverage [%] | # Peptides | # PSMs | # Unique Peptides | # AAs | MW [kDa] | calc. pI |
|---|---|---|---|---|---|---|---|---|---|---|---|---|---|
| High | Master Protein | A0PJE6 | Pccb protein (Fragment) OS=Mus musculus OX=10090 GN=Pccb PE=2 SV=1 | FAUX | Mouse | 166,707 | 65 | 24 | 305 | 2 | 539 | 58,1 | 7,66 |
| High | Master Protein | Q08EK5 | Keratin 77 OS=Mus musculus OX=10090 GN=Krt77 PE=1 SV=1 | FAUX | Mouse | 82,021 | 14 | 13 | 354 | 3 | 572 | 61,3 | 8,02 |
| High | Master Protein | A2A513 | Keratin 10 OS=Mus musculus OX=10090 GN=Krt10 PE=1 SV=1 | FAUX | Mouse | 79,167 | 15 | 12 | 352 | 1 | 561 | 57 | 5,07 |
| High | Master Protein | D3YYT1 | Cytokine-like nuclear factor N-PAC OS=Mus musculus OX=10090 GN=Glyr1 PE=1 SV=1 | FAUX | Mouse | 62,163 | 33 | 13 | 45 | 13 | 552 | 60,4 | 9,13 |
| High | Master Protein | Q9CPQ9 | Fructose-bisphosphate aldolase OS=Mus musculus OX=10090 GN=Aldoart1 PE=1 SV=1 | FAUX | Mouse | 56,019 | 35 | 12 | 151 | 1 | 364 | 39,3 | 7,37 |
| High | Master Protein | Q3SYP5 | Keratin 16 OS=Mus musculus OX=10090 GN=Krt16 PE=1 SV=1 | FAUX | Mouse | 54,559 | 18 | 8 | 144 | 5 | 469 | 51,6 | 5,2 |
| High | Master Protein | Q3TT11 | Uncharacterized protein OS=Mus musculus OX=10090 GN=Hsd17b4 PE=2 SV=1 | FAUX | Mouse | 50,701 | 22 | 12 | 81 | 12 | 735 | 79,4 | 8,57 |
| High | Master Protein | Q5NC80 | Nucleoside diphosphate kinase (Fragment) OS=Mus musculus OX=10090 GN=Nme1 PE=1 SV=1 | FAUX | Mouse | 29,609 | 61 | 6 | 37 | 1 | 127 | 14,1 | 9,09 |
| High | Master Protein | A0A8J2HFW6 | Mitochondrial (Mus musculus) OS=Cotesia congregata OX=51543 GN=HICCMSTLAB_LOCUS7100 PE=3 SV=1 | FAUX | Mouse | 26,89 | 10 | 6 | 40 | 5 | 767 | 83,6 | 8,38 |
| High | Master Protein | H3BKR2 | Guanine nucleotide binding protein (G protein), beta 1 (Fragment) OS=Mus musculus OX=10090 GN=Gnb1 PE=1 SV=8 | FAUX | Mouse | 24,799 | 26 | 6 | 61 | 2 | 273 | 30,2 | 6,21 |
| High | Master Protein | Q80V08 | Large ribosomal subunit protein uL22 (Fragment) OS=Mus musculus OX=10090 GN=Rpl17 PE=2 SV=1 | FAUX | Mouse | 22,527 | 26 | 4 | 27 | 4 | 194 | 22,4 | 10,33 |
| High | Master Protein | B1AQ78 | Keratin 19 OS=Mus musculus OX=10090 GN=Krt19 PE=1 SV=1 | FAUX | Mouse | 20,914 | 11 | 6 | 175 | 3 | 403 | 44,5 | 5,39 |
| High | Master Protein | Q9JJ20 | 14-3-3 protein sigma OS=Mus musculus OX=10090 GN=Sfn PE=2 SV=1 | FAUX | Mouse | 17,576 | 15 | 4 | 59 | 1 | 248 | 27,8 | 4,78 |
| High | Master Protein | Q3U292 | H15 domain-containing protein OS=Mus musculus OX=10090 GN=H1f3 PE=2 SV=1 | FAUX | Mouse | 15,344 | 22 | 4 | 34 | 1 | 221 | 22,1 | 11,03 |
| High | Master Protein | A0A0A6YW87 | Cornichon family AMPA receptor auxiliary protein 4 OS=Mus musculus OX=10090 GN=Cnih4 PE=1 SV=1 | FAUX | Mouse | 14,166 | 26 | 1 | 26 | 1 | 78 | 9,2 | 8,21 |
| High | Master Protein | B2RXT3 | oxoglutarate dehydrogenase (succinyl-transferring) OS=Mus musculus OX=10090 GN=Ogdhl PE=1 SV=1 | FAUX | Mouse | 13,375 | 5 | 4 | 8 | 2 | 1010 | 114,5 | 6,84 |
| High | Master Protein | F6V084 | Thioredoxin-related transmembrane protein 1 (Fragment) OS=Mus musculus OX=10090 GN=Tmx1 PE=1 SV=1 | FAUX | Mouse | 12,061 | 18 | 2 | 11 | 2 | 124 | 14,4 | 5,06 |
| High | Master Protein | Q543F3 | Calponin OS=Mus musculus OX=10090 GN=Cnn2 PE=1 SV=1 | FAUX | Mouse | 8,844 | 9 | 2 | 7 | 1 | 305 | 33,1 | 7,62 |

| | | | | | | | | | | | | | |
|---|---|---|---|---|---|---|---|---|---|---|---|---|---|
| High | Master Protein | Q3UGB5 | DAZ associated protein 1 OS=Mus musculus OX=10090 GN=Dazap1 PE=1 SV=1 | FAUX | Mouse | 8,403 | 8 | 2 | 3 | 2 | 406 | 43,1 | 8,56 |
| High | Master Protein | A0A0R4J293 | Transglutaminase 1, K polypeptide OS=Mus musculus OX=10090 GN=Tgm1 PE=1 SV=1 | FAUX | Mouse | 8,228 | 4 | 3 | 5 | 3 | 815 | 89,7 | 6,58 |
| High | Master Protein | A0A077S9N1 | Lysozyme f3 OS=Mus musculus OX=10090 GN=Lyz1 PE=3 SV=1 | VRAI | Mouse | 7,618 | 8 | 1 | 23 | 1 | 148 | 16,8 | 9,41 |
| High | Master Protein | A0A0R4J1N8 | Retinol dehydrogenase 13 (all-trans and 9-cis) OS=Mus musculus OX=10090 GN=Rdh13 PE=1 SV=1 | FAUX | Mouse | 7,026 | 10 | 2 | 2 | 2 | 299 | 32,3 | 9,52 |
| High | Master Protein | B9EI85 | Histone H3 OS=Mus musculus OX=10090 GN=Hist2h3b PE=2 SV=1 | FAUX | Mouse | 6,075 | 35 | 3 | 4 | 3 | 136 | 15,4 | 11,27 |
| High | Master Protein | Q9CV17 | IF rod domain-containing protein (Fragment) OS=Mus musculus OX=10090 GN=Krt24 PE=2 SV=1 | FAUX | Mouse | 6,043 | 8 | 2 | 65 | 1 | 304 | 33,9 | 4,73 |
| High | Master Protein | Q571M0 | MKIAA4032 protein (Fragment) OS=Mus musculus OX=10090 GN=Mfn1 PE=2 SV=1 | FAUX | Mouse | 5,346 | 2 | 1 | 2 | 1 | 749 | 84,5 | 6,65 |
| High | Master Protein | D6RGM4 | Sphingomyelin phosphodiesterase 2, neutral OS=Mus musculus OX=10090 GN=Smpd2 PE=1 SV=1 | FAUX | Mouse | 4,832 | 14 | 1 | 5 | 1 | 83 | 10,1 | 9,25 |
| High | Master Protein | Q6S390 | Plectin 4 OS=Mus musculus OX=10090 GN=Plec PE=2 SV=1 | FAUX | Mouse | 4,615 | 0 | 2 | 5 | 2 | 4449 | 506,1 | 5,71 |
| High | Master Protein | Q9CVL8 | Signal recognition particle receptor alpha subunit N-terminal domain-containing protein (Fragment) OS=Mus musculus OX=10090 GN=Srpr PE=2 SV=1 | FAUX | Mouse | 4,37 | 6 | 1 | 2 | 1 | 161 | 18,6 | 9,23 |
| High | Master Protein | A0A1D5RM62 | Hydroxysteroid dehydrogenase like 1 (Fragment) OS=Mus musculus OX=10090 GN=Hsdl1 PE=1 SV=1 | FAUX | Mouse | 4,287 | 13 | 1 | 1 | 1 | 101 | 11,2 | 7,43 |
| High | Master Protein | Q3U844 | Protein kinase domain-containing protein OS=Mus musculus OX=10090 GN=Csnk1a1 PE=2 SV=1 | FAUX | Mouse | 4,004 | 3 | 1 | 1 | 1 | 325 | 37,5 | 9,48 |
| High | Master Protein | Q497I3 | Fatty acid binding protein 5, epidermal OS=Mus musculus OX=10090 GN=Fabp5 PE=1 SV=1 | FAUX | Mouse | 3,894 | 7 | 1 | 10 | 1 | 135 | 15,1 | 6,54 |
| High | Master Protein | Q3UBU1 | Dihydroxyacetone phosphate acyltransferase OS=Mus musculus OX=10090 GN=Gnpat PE=2 SV=1 | FAUX | Mouse | 3,705 | 2 | 1 | 1 | 1 | 678 | 76,8 | 8,1 |
| High | Master Protein | D3Z6W2 | Tyrosine-protein phosphatase non-receptor type OS=Mus musculus OX=10090 GN=Ptpn2 PE=1 SV=1 | FAUX | Mouse | 3,647 | 5 | 1 | 1 | 1 | 363 | 42,3 | 8,29 |
| High | Master Protein | A2AW86 | Lymphocyte antigen 75 OS=Mus musculus OX=10090 GN=Ly75 PE=1 SV=1 | FAUX | Mouse | 3,178 | 1 | 1 | 1 | 1 | 1723 | 197,3 | 6,39 |
| High | Master Protein | Q69ZW6 | MKIAA0889 protein (Fragment) OS=Mus musculus OX=10090 GN=Soga1 PE=2 SV=1 | FAUX | Mouse | 3,143 | 3 | 1 | 5 | 1 | 830 | 93 | 8,38 |
| High | Master Protein | A0A087WQM9 | Serine (or cysteine) peptidase inhibitor, clade B, member 5 (Fragment) OS=Mus musculus OX=10090 GN=Serpinb5 PE=1 SV=1 | FAUX | Mouse | 3,092 | 12 | 1 | 2 | 1 | 74 | 8,5 | 5,14 |
| High | Master Protein | A0A0A6YX96 | Folylpolyglutamyl synthetase OS=Mus musculus OX=10090 GN=Fpgs PE=1 SV=1 | FAUX | Mouse | 3,052 | 8 | 1 | 1 | 1 | 189 | 21,5 | 9,16 |
| High | Master Protein | B2FDE8 | IBA57 homolog, iron-sulfur cluster assembly (Fragment) OS=Mus musculus OX=10090 GN=Iba57 PE=1 SV=1 | FAUX | Mouse | 3,044 | 6 | 1 | 1 | 1 | 215 | 22,6 | 9,57 |
| High | Master Protein | A0A0U1RPH8 | Kinesin family member 22 (Fragment) OS=Mus musculus OX=10090 GN=Kif22 PE=1 SV=1 | FAUX | Mouse | 2,874 | 10 | 1 | 1 | 1 | 161 | 17,8 | 4,79 |
| High | Master Protein | Z4YJD9 | Keratin 12 OS=Mus musculus OX=10090 GN=Krt12 PE=1 SV=1 | FAUX | Mouse | 2,865 | 2 | 1 | 1 | 1 | 487 | 52,5 | 4,84 |
| High | Master Protein | Q6A064 | MKIAA0472 protein (Fragment) OS=Mus musculus OX=10090 GN=Dstyk PE=2 SV=1 | FAUX | Mouse | 2,83 | 1 | 1 | 5 | 1 | 658 | 75,2 | 7,42 |
| High | Master Protein | D3Z0S9 | Tumor susceptibility gene 101 (Fragment) OS=Mus musculus OX=10090 GN=Tsg101 PE=1 SV=2 | FAUX | Mouse | 2,822 | 7 | 1 | 1 | 1 | 145 | 15,5 | 6,77 |
| High | Master Protein | D3YXK0 | Janus kinase and microtubule interacting protein 2 OS=Mus musculus OX=10090 GN=Jakmip2 PE=1 SV=2 | FAUX | Mouse | 2,736 | 1 | 1 | 6 | 1 | 820 | 96,3 | 5,97 |
| High | Master Protein | Q45FA7 | Small conductance calcium-activated potassium channel 3 OS=Mus musculus OX=10090 GN=Kcnn3 PE=2 SV=1 | FAUX | Mouse | 2,72 | 4 | 1 | 1 | 1 | 731 | 81,1 | 8,95 |
| High | Master Protein | Q5SUD5 | SCO1 cytochrome c oxidase assembly protein OS=Mus musculus OX=10090 GN=Sco1 PE=1 SV=1 | FAUX | Mouse | 2,473 | 3 | 1 | 1 | 1 | 289 | 32,1 | 8,06 |
| High | Master Protein | Q5SWJ8 | Protein transport protein Sec61 subunit gamma OS=Mus musculus OX=10090 GN=Sec61g PE=2 SV=1 | FAUX | Mouse | 2,366 | 19 | 1 | 1 | 1 | 68 | 7,7 | 9,99 |
| High | Master Protein | Q3U6T2 | Alpha-galactosidase OS=Mus musculus OX=10090 GN=Naga PE=2 SV=1 | FAUX | Mouse | 2,269 | 3 | 1 | 3 | 1 | 415 | 47,2 | 6,44 |
| High | Master Protein | A0A0G2JFW6 | Regulation of nuclear pre-mRNA domain containing 2 (Fragment) OS=Mus musculus OX=10090 GN=Rprd2 PE=1 SV=1 | FAUX | Mouse | 2,184 | 1 | 1 | 2 | 1 | 1386 | 147,6 | 6,93 |
| High | Master Protein | Q5SQ20 | Pescadillo homolog OS=Mus musculus OX=10090 GN=Pes1 PE=1 SV=1 | FAUX | Mouse | 2,132 | 2 | 1 | 1 | 1 | 588 | 68,2 | 6,84 |
| High | Master Protein | P62843 | 40S ribosomal protein S15 OS=Mus musculus OX=10090 GN=Rps15 PE=1 SV=2 | FAUX | Mouse | 1,544 | 8 | 1 | 1 | 1 | 145 | 17 | 10,39 |
| High | Master Protein | P99027 | 60S acidic ribosomal protein P2 OS=Mus musculus OX=10090 GN=Rplp2 PE=1 SV=3 | FAUX | Mouse | 5,454 | 14 | 1 | 1 | 1 | 115 | 11,6 | 4,54 |
| High | Master Protein | Q3TUI6 | ABC_transp_aux domain-containing protein OS=Mus musculus OX=10090 GN=Ift52 PE=2 SV=1 | FAUX | Mouse | 2,677 | 4 | 1 | 1 | 1 | 271 | 30,2 | 5,29 |
| High | Master Protein | Q9JM76 | Actin-related protein 2/3 complex subunit 3 OS=Mus musculus OX=10090 GN=Arpc3 PE=1 SV=3 | FAUX | Mouse | 2,754 | 8 | 1 | 2 | 1 | 178 | 20,5 | 8,59 |
| High | Master Protein | A2ATT5 | ADP-sugar pyrophosphatase (Fragment) OS=Mus musculus OX=10090 GN=Nudt5 PE=1 SV=1 | FAUX | Mouse | 1,212 | 8 | 1 | 1 | 1 | 161 | 17,7 | 5,4 |
| High | Master Protein | A1A4T2 | Alpha glucosidase 2 alpha neutral subunit OS=Mus musculus OX=10090 GN=Ganab PE=1 SV=1 | FAUX | Mouse | 1,444 | 1 | 1 | 1 | 1 | 966 | 109,3 | 6,15 |
| High | Master Protein | Q8K298 | Anillin OS=Mus musculus OX=10090 GN=Anln PE=1 SV=2 | FAUX | Mouse | 1,305 | 1 | 1 | 1 | 1 | 1121 | 122,7 | 6,98 |
| High | Master Protein | A0A087WNV1 | Arf-GAP domain and FG repeat-containing protein 1 OS=Mus musculus OX=10090 GN=Agfg1 PE=1 SV=1 | FAUX | Mouse | 1,625 | 2 | 1 | 1 | 1 | 521 | 53,9 | 8,92 |
| High | Master Protein | Q5EBI8 | ATP synthase subunit e, mitochondrial OS=Mus musculus OX=10090 GN=Atp5k PE=2 SV=1 | FAUX | Mouse | 2,727 | 17 | 1 | 1 | 1 | 71 | 8,2 | 8,44 |
| High | Master Protein | A6H6S8 | BCL2-associated athanogene 4 OS=Mus musculus OX=10090 GN=Bag4 PE=1 SV=1 | FAUX | Mouse | 7,049 | 4 | 1 | 1 | 1 | 457 | 49,1 | 5,34 |
| High | Master Protein | Q3TYE1 | Casein kinase I isoform epsilon OS=Mus musculus OX=10090 GN=Csnk1e PE=1 SV=1 | FAUX | Mouse | 3,861 | 2 | 1 | 1 | 1 | 368 | 42,2 | 8,9 |
| High | Master Protein | Q60737 | Casein kinase II subunit alpha OS=Mus musculus OX=10090 GN=Csnk2a1 PE=1 SV=2 | FAUX | Mouse | 6,402 | 4 | 1 | 1 | 1 | 391 | 45,1 | 7,74 |
| High | Master Protein | Q3UCW4 | Cathepsin D OS=Mus musculus OX=10090 GN=Ctsd PE=2 SV=1 | FAUX | Mouse | 6,022 | 7 | 3 | 3 | 3 | 410 | 44,9 | 7,15 |
| High | Master Protein | A0A0U1RP13 | CDP-diacylglycerol--inositol 3-phosphatidyltransferase OS=Mus musculus OX=10090 GN=Cdipt PE=1 SV=1 | FAUX | Mouse | 1,788 | 5 | 1 | 1 | 1 | 185 | 20,3 | 8,48 |
| High | Master Protein | Q5SUH7 | Clathrin interactor 1 OS=Mus musculus OX=10090 GN=Clint1 PE=1 SV=1 | FAUX | Mouse | 20,446 | 8 | 4 | 4 | 4 | 623 | 67,7 | 6,42 |
| High | Master Protein | H3BKW0 | Cleavage and polyadenylation-specificity factor subunit 6 (Fragment) OS=Mus musculus OX=10090 GN=Cpsf6 PE=1 SV=1 | FAUX | Mouse | 3,599 | 14 | 1 | 1 | 1 | 182 | 21,2 | 9,73 |
| High | Master Protein | Q3UIT2 | COP9 signalosome complex subunit 6 OS=Mus musculus OX=10090 GN=Cops6 PE=1 SV=1 | FAUX | Mouse | 1,993 | 2 | 1 | 1 | 1 | 324 | 35,9 | 5,73 |
| High | Master Protein | Q54AC6 | Cox17p OS=Mus musculus OX=10090 GN=COX17 PE=1 SV=1 | FAUX | Mouse | 6,175 | 27 | 1 | 1 | 1 | 63 | 6,8 | 7,69 |
| High | Master Protein | A2A432 | Cullin-4B OS=Mus musculus OX=10090 GN=Cul4b PE=1 SV=1 | FAUX | Mouse | 3,51 | 1 | 1 | 1 | 1 | 970 | 110,6 | 8,37 |
| High | Master Protein | Q3UMI9 | Cullin-associated NEDD8-dissociated protein 1 (Fragment) OS=Mus musculus OX=10090 GN=Cand1 PE=2 SV=1 | FAUX | Mouse | 1,455 | 1 | 1 | 1 | 1 | 700 | 77,9 | 5,54 |
| High | Master Protein | P97315 | Cysteine and glycine-rich protein 1 OS=Mus musculus OX=10090 GN=Csrp1 PE=1 SV=3 | FAUX | Mouse | 3,443 | 8 | 1 | 1 | 1 | 193 | 20,6 | 8,57 |
| High | Master Protein | P0CG14 | Decreased expression in renal and prostate cancer protein OS=Mus musculus OX=10090 GN=Derpc PE=1 SV=1 | FAUX | Mouse | 1,14 | 2 | 1 | 1 | 1 | 533 | 52,2 | 12,32 |
| High | Master Protein | Q3U9G9 | Delta(14)-sterol reductase LBR OS=Mus musculus OX=10090 GN=Lbr PE=1 SV=2 | FAUX | Mouse | 1,42 | 1 | 1 | 1 | 1 | 626 | 71,4 | 9,36 |
| High | Master Protein | Q8BHC4 | Dephospho-CoA kinase domain-containing protein OS=Mus musculus OX=10090 GN=Dcakd PE=1 SV=1 | FAUX | Mouse | 2,046 | 6 | 1 | 1 | 1 | 231 | 26,5 | 9,58 |
| High | Master Protein | A0A494BBJ6 | DNA damage-binding protein 1 (Fragment) OS=Mus musculus OX=10090 GN=Ddb1 PE=1 SV=1 | FAUX | Mouse | 4,068 | 7 | 1 | 1 | 1 | 196 | 21,4 | 5,16 |
| High | Master Protein | A0A3B2WBC6 | DNA-directed RNA polymerase II subunit GRINL1A OS=Mus musculus OX=10090 GN=Polr2m PE=4 SV=1 | FAUX | Mouse | 9,593 | 29 | 1 | 1 | 1 | 84 | 8,6 | 10,48 |
| High | Master Protein | K3W4M1 | DnaJ homolog subfamily C member 4 OS=Mus musculus OX=10090 GN=Dnajc4 PE=4 SV=1 | FAUX | Mouse | 1,257 | 3 | 1 | 1 | 1 | 261 | 30 | 9,89 |
| High | Master Protein | Q3U505 | Dolichyl-diphosphooligosaccharide--protein glycosyltransferase subunit 2 (Fragment) OS=Mus musculus OX=10090 GN=Rpn2 PE=2 SV=1 | FAUX | Mouse | 23,071 | 10 | 4 | 4 | 4 | 659 | 72,1 | 6,49 |
| High | Master Protein | Q6PDX6 | E3 ubiquitin-protein ligase Rnf220 OS=Mus musculus OX=10090 GN=Rnf220 PE=1 SV=1 | FAUX | Mouse | 12,396 | 11 | 4 | 4 | 4 | 566 | 62,7 | 6,04 |
| High | Master Protein | P83940 | Elongin-C OS=Mus musculus OX=10090 GN=Eloc PE=1 SV=1 | FAUX | Mouse | 1,402 | 9 | 1 | 1 | 1 | 112 | 12,5 | 4,78 |
| High | Master Protein | Q9DCS3 | Enoyl-[acyl-carrier-protein] reductase, mitochondrial OS=Mus musculus OX=10090 GN=Mecr PE=1 SV=2 | FAUX | Mouse | 5,046 | 3 | 1 | 1 | 1 | 373 | 40,3 | 9,07 |
| High | Master Protein | Q811B2 | Epb4.1l2 protein OS=Mus musculus OX=10090 GN=Epb41l2 PE=2 SV=1 | FAUX | Mouse | 1,406 | 1 | 1 | 1 | 1 | 689 | 76,9 | 6,61 |
| High | Master Protein | Q8JZQ9 | Eukaryotic translation initiation factor 3 subunit B OS=Mus musculus OX=10090 GN=Eif3b PE=1 SV=1 | FAUX | Mouse | 5,751 | 3 | 1 | 1 | 1 | 803 | 91,3 | 5,02 |
| High | Master Protein | Q9Z1D1 | Eukaryotic translation initiation factor 3 subunit G OS=Mus musculus OX=10090 GN=Eif3g PE=1 SV=2 | FAUX | Mouse | 1,99 | 3 | 1 | 1 | 1 | 320 | 35,6 | 5,9 |
| High | Master Protein | A2AJ71 | Far upstream element (FUSE)-binding protein 3 (Fragment) OS=Mus musculus OX=10090 GN=Fubp3 PE=1 SV=1 | FAUX | Mouse | 32,55 | 69 | 5 | 13 | 1 | 97 | 10,4 | 6,79 |
| High | Master Protein | A0A286YDK2 | Growth hormone-inducible transmembrane protein (Fragment) OS=Mus musculus OX=10090 GN=Ghitm PE=1 SV=1 | FAUX | Mouse | 1,391 | 5 | 1 | 1 | 1 | 198 | 21,7 | 9,91 |
| High | Master Protein | Q3UXX9 | HCLS1-associated protein X-1 OS=Mus musculus OX=10090 GN=Hax1 PE=2 SV=1 | FAUX | Mouse | 1,656 | 6 | 1 | 1 | 1 | 196 | 22,2 | 4,74 |
| High | Master Protein | A2VCT7 | Hexosyltransferase (Fragment) OS=Mus musculus OX=10090 GN=Chsy1 PE=2 SV=1 | FAUX | Mouse | 1,089 | 1 | 1 | 1 | 1 | 747 | 85,7 | 8,92 |
| High | Master Protein | A0A1B0GQX9 | High mobility group protein B2 (Fragment) OS=Mus musculus OX=10090 GN=Hmgb2 PE=1 SV=1 | FAUX | Mouse | 1,464 | 8 | 1 | 1 | 1 | 99 | 11,5 | 9,89 |
| High | Master Protein | Q61081 | Hsp90 co-chaperone Cdc37 OS=Mus musculus OX=10090 GN=Cdc37 PE=1 SV=1 | FAUX | Mouse | 5,475 | 4 | 1 | 1 | 1 | 379 | 44,6 | 5,34 |
| High | Master Protein | Q8BK58 | HSPB1-associated protein 1 OS=Mus musculus OX=10090 GN=Hspbap1 PE=1 SV=2 | FAUX | Mouse | 1,546 | 2 | 1 | 1 | 1 | 483 | 54,4 | 5,27 |
| High | Master Protein | P00493 | Hypoxanthine-guanine phosphoribosyltransferase OS=Mus musculus OX=10090 GN=Hprt1 PE=1 SV=3 | FAUX | Mouse | 3,492 | 6 | 1 | 1 | 1 | 218 | 24,6 | 6,68 |
| High | Master Protein | Q3TGZ3 | Isocitrate dehydrogenase [NAD] subunit, mitochondrial OS=Mus musculus OX=10090 GN=Idh3g PE=1 SV=1 | FAUX | Mouse | 5,479 | 6 | 1 | 1 | 1 | 393 | 42,8 | 9,01 |
| High | Master Protein | Q3UKA1 | J domain-containing protein OS=Mus musculus OX=10090 GN=Dnajc3 PE=2 SV=1 | FAUX | Mouse | 2,366 | 2 | 1 | 1 | 1 | 504 | 57,5 | 5,95 |
| High | Master Protein | G3X961 | Kelch domain-containing protein 4 OS=Mus musculus OX=10090 GN=Klhdc4 PE=1 SV=1 | FAUX | Mouse | 3,287 | 2 | 1 | 1 | 1 | 584 | 64,8 | 5,66 |
| High | Master Protein | A0A1B0GSR7 | Keratin, type I cytoskeletal 10 OS=Mus musculus OX=10090 GN=Krt10 PE=1 SV=1 | FAUX | Mouse | 1,273 | 10 | 1 | 1 | 1 | 97 | 9,3 | 8,48 |
| High | Master Protein | A1L317 | Keratin, type I cytoskeletal 24 OS=Mus musculus OX=10090 GN=Krt24 PE=2 SV=2 | FAUX | Mouse | 2,076 | 3 | 1 | 1 | 1 | 512 | 54 | 5,01 |
| High | Master Protein | Q3UR33 | Lamina-associated polypeptide 2 OS=Mus musculus OX=10090 GN=Tmpo PE=2 SV=1 | FAUX | Mouse | 3,361 | 2 | 1 | 1 | 1 | 452 | 50,3 | 9,41 |
| High | Master Protein | Q6PB66 | Leucine-rich PPR motif-containing protein, mitochondrial OS=Mus musculus OX=10090 GN=Lrpprc PE=1 SV=2 | FAUX | Mouse | 3,489 | 1 | 1 | 1 | 1 | 1392 | 156,5 | 6,83 |
| High | Master Protein | Q8R1F0 | Leydig cell tumor 10 kDa protein homolog OS=Mus musculus OX=10090 GN=D8Ertd738e PE=3 SV=1 | FAUX | Mouse | 2,86 | 16 | 2 | 2 | 2 | 94 | 10,2 | 11,63 |
| High | Master Protein | Q8CFK5 | Lysine--tRNA ligase OS=Mus musculus OX=10090 GN=Kars PE=2 SV=1 | FAUX | Mouse | 1,311 | 2 | 1 | 1 | 1 | 622 | 71 | 6,43 |
| High | Master Protein | Q9D880 | Mitochondrial import inner membrane translocase subunit TIM50 OS=Mus musculus OX=10090 GN=Timm50 PE=1 SV=1 | FAUX | Mouse | 1,727 | 3 | 1 | 1 | 1 | 353 | 39,8 | 8,13 |
| High | Master Protein | Q2TAW4 | Myosin, heavy polypeptide 6, cardiac muscle, alpha OS=Mus musculus OX=10090 GN=Myh6 PE=2 SV=1 | FAUX | Mouse | 5,138 | 1 | 1 | 1 | 1 | 1938 | 223,4 | 5,73 |
| High | Master Protein | Q9CQ75 | NADH dehydrogenase [ubiquinone] 1 alpha subcomplex subunit 2 OS=Mus musculus OX=10090 GN=Ndufa2 PE=1 SV=3 | FAUX | Mouse | 5,103 | 14 | 1 | 1 | 1 | 99 | 10,9 | 9,99 |
| High | Master Protein | Q9CPP6 | NADH dehydrogenase [ubiquinone] 1 alpha subcomplex subunit 5 OS=Mus musculus OX=10090 GN=Ndufa5 PE=1 SV=3 | FAUX | Mouse | 1,688 | 8 | 1 | 1 | 1 | 116 | 13,4 | 8,1 |

| | | | | | | | | | | | | | |
|---|---|---|---|---|---|---|---|---|---|---|---|---|---|
| High | Master Protein | F7BIJ5 | Nuclear receptor-interacting protein 2 (Fragment) OS=Mus musculus OX=10090 GN=Nrip2 PE=4 SV=1 | FAUX | Mouse | 1,359 | 8 | 1 | 1 | 1 | 113 | 12,1 | 8,16 |
| High | Master Protein | Q3TGU7 | Peptidase_M24 domain-containing protein OS=Mus musculus OX=10090 GN=Pa2g4 PE=1 SV=1 | FAUX | Mouse | 1,626 | 2 | 1 | 1 | 1 | 394 | 43,7 | 6,86 |
| High | Master Protein | Q9D3J1 | Peptidyl-prolyl cis-trans isomerase OS=Mus musculus OX=10090 GN=Ppil4 PE=2 SV=1 | FAUX | Mouse | 1,687 | 2 | 1 | 1 | 1 | 460 | 53,2 | 5,43 |
| High | Master Protein | Q3ULN5 | peptidylprolyl isomerase OS=Mus musculus OX=10090 GN=Fkbp1a PE=2 SV=1 | FAUX | Mouse | 6,253 | 13 | 1 | 1 | 1 | 108 | 12 | 8,16 |
| High | Master Protein | Q6GT24 | Peroxiredoxin-6 OS=Mus musculus OX=10090 GN=Prdx6 PE=1 SV=1 | FAUX | Mouse | 2,676 | 4 | 1 | 1 | 1 | 224 | 24,8 | 6,37 |
| High | Master Protein | A0A494BAS6 | Peroxisomal 2,4-dienoyl-CoA reductase [(3E)-enoyl-CoA-producing] OS=Mus musculus OX=10090 GN=Decr2 PE=1 SV=1 | FAUX | Mouse | 3,67 | 4 | 1 | 1 | 1 | 268 | 28,8 | 9,88 |
| High | Master Protein | Q8CD98 | PFK domain-containing protein OS=Mus musculus OX=10090 GN=Pfkl PE=2 SV=1 | FAUX | Mouse | 1,118 | 5 | 1 | 1 | 1 | 465 | 51,7 | 7,72 |
| High | Master Protein | Q9EP69 | Phosphatidylinositol-3-phosphatase SAC1 OS=Mus musculus OX=10090 GN=Sacm1l PE=1 SV=1 | FAUX | Mouse | 2,509 | 3 | 2 | 2 | 2 | 587 | 66,9 | 7,3 |
| High | Master Protein | Q3UEB3 | Poly(U)-binding-splicing factor PUF60 OS=Mus musculus OX=10090 GN=Puf60 PE=1 SV=2 | FAUX | Mouse | 1,202 | 1 | 1 | 1 | 1 | 564 | 60,2 | 5,29 |
| High | Master Protein | F6SQH7 | Polymerase delta-interacting protein 2 (Fragment) OS=Mus musculus OX=10090 GN=Poldip2 PE=1 SV=1 | FAUX | Mouse | 2,282 | 4 | 1 | 1 | 1 | 284 | 32,6 | 7,91 |
| High | Master Protein | A0A2R8VHP3 | Predicted pseudogene 5478 OS=Mus musculus OX=10090 GN=Gm5478 PE=1 SV=1 | FAUX | Mouse | 25,172 | 10 | 7 | 27 | 1 | 535 | 57,9 | 6,2 |
| High | Master Protein | P42208 | Septin-2 OS=Mus musculus OX=10090 GN=Septin2 PE=1 SV=2 | FAUX | Mouse | 1,077 | 2 | 1 | 1 | 1 | 361 | 41,5 | 6,55 |
